# The Lyme Disease agent co-opts adiponectin receptor-mediated signaling in its arthropod vector

**DOI:** 10.1101/2021.09.13.460039

**Authors:** Xiaotian Tang, Yongguo Cao, Gunjan Arora, Jesse Hwang, Andaleeb Sajid, Courtney L. Brown, Sameet Mehta, Alejandro Marín-López, Yu-Min Chuang, Ming-Jie Wu, Hongwei Ma, Utpal Pal, Sukanya Narasimhan, Erol Fikrig

**Affiliations:** Section of Infectious Diseases, Department of Internal Medicine, School of Medicine, Yale University, New Haven, CT, USA; Department of Clinical Veterinary Medicine, and Key Laboratory for Zoonosis Research, Ministry of Education, College of Veterinary Medicine, Jilin University, Changchun, China; Yale Combined Program in the Biological and Biomedical Sciences, Yale University, New Haven, CT, USA; Yale Center for Genome Analysis, Yale University, New Haven, CT, USA; Department of Microbiology, School of Basic Medicine, Fourth Military Medical University, Xi’an, Shaanxi, China; Department of Veterinary Medicine, University of Maryland, College Park, Maryland, USA

**Author notes:** Corresponding authors: Erol Fikrig, Section of Infectious Diseases, Department of Internal Medicine, Yale University School of Medicine,; Xiaotian Tang. These authors contributed equally to the manuscript (Xiaotian Tang, Yongguo Cao).

**Keywords:** Adiponectin receptor, *Ixodes scapularis*, *Borrelia burgdorferi*, Phospholipid metabolism, C1q-like protein 3 protein

## Abstract

Adiponectin-mediated pathways contribute to mammalian homeostasis; however, little is known about adiponectin and adiponectin receptor signaling in arthropods. In this study, we demonstrate that *Ixodes scapularis* ticks have an adiponectin receptor-like protein (ISARL) but lack adiponectin – suggesting activation by alternative pathways. *ISARL* expression is significantly upregulated in the tick gut after *Borrelia burgdorferi* infection suggesting that ISARL-signaling may be co-opted by the Lyme disease agent. Consistent with this, RNA interference (RNAi)-mediated silencing of *ISARL* significantly reduced the *B. burgdorferi* burden in the tick. RNA-seq-based transcriptomics and RNAi assays demonstrate that ISARL-mediated phospholipid metabolism by phosphatidylserine synthase I is associated with *B. burgdorferi* survival. Furthermore, the tick complement C1q-like protein 3 interacts with ISARL, and *B. burgdorferi* facilitates this process. This study identifies a new tick metabolic pathway that is connected to the life cycle of the Lyme disease spirochete.

**Significance Statement:** Adiponectin binds to adiponectin receptors and participates in glucose and lipid metabolism in mammals. In this study, we found that ticks have an adiponectin receptor-like protein but lack adiponectin. Importantly, we demonstrated that the Lyme disease agent, *Borrelia burgdorferi*, takes advantage of the adiponectin receptor signaling pathway to establish infection in its arthropod vector, *Ixodes scapularis*. Our study sheds light on the understanding of *Borrelia*-tick interactions and provides insights into how a human infectious disease agent may evolve to manipulate host metabolism for its own benefits. Understanding this pathway may lead to new ways to interfere with the *Borrelia* life cycle, and this mechanism may be applicable to additional microbes that are transmitted by ticks, mosquitoes or other arthropods.

## Introduction

Adiponectin, adipocyte complement related protein of 30 kDa (or Acrp30), plays important roles in the regulation of metabolism, insulin sensitivity, and inflammation across species (Kadowaki et al., 2006; Ouchi and Walsh, 2007; Yamauchi et al., 2002). Adiponectin mediates its actions mainly via binding adiponectin receptors with its globular C1q domain (Buechler et al., 2010; Yamauchi et al., 2002). Two adiponectin receptors, AdipoR1 and AdipoR2, have been identified in mammals (Yamauchi et al., 2003). AdipoR1 and R2 belong to a family of membrane receptors predicted to contain seven transmembrane domains with an internal N terminus and an external C terminus (Yamauchi et al., 2003). AdipoR1 has a higher binding affinity for the globular form of adiponectin, whereas AdipoR2 has a greater affinity for full length adiponectin (Yamauchi et al., 2003). Interestingly, AdipoR1 and AdipoR2 double-knockout mice have increased triglyceride levels, and exhibit insulin resistance, demonstrating that AdipoR1 and AdipoR2 regulate lipid and glucose homeostasis (Kadowaki et al., 2006; Yamauchi et al., 2007). In yeast, a homolog of mammalian adiponectin receptors, ORE20/PHO36, is involved in lipid and phosphate metabolism (Karpichev et al., 2002). PHO36, can also interact with a plant protein, osmotin, a homolog of mammalian adiponectin, thereby controlling apoptosis in yeast (Narasimhan et al., 2005). Adiponectin and adiponectin receptors in disease-transmitting arthropods have not been characterized. By utilizing the amino acid sequence homology search in other model arthropods, adiponectin was not identified from *Drosophila melanogaster*, however, an adiponectin receptor which regulates insulin secretion and controls glucose and lipid metabolism was characterized (Kwak et al., 2013). In addition, Zhu et al. (2008) cloned an adiponectin-like receptor gene from the silk moth, *Bombyx mori*, and found that infection with *B. mori* nucleopolyhedrovirus significantly increased adiponectin receptor mRNA levels in the midgut of susceptible *B. mori*, suggesting an association with pathogen infectivity.

*Ixodes scapularis*, the black-legged tick, is an important vector of the Lyme disease agent, *Borrelia burgdorferi* (Estrada-Peña and Jongejan, 1999), which causes approximately 300,000 cases annually in the United States (Rosenberg et al., 2018). *B. burgdorferi* is acquired when larval or nymphal ticks feed on infected animals, and is transmitted by nymphs or adults to vertebrate hosts (Kurokawa et al., 2020). Lyme disease in humans manifests as a multisystem disorder of the skin and other organs (e.g., joints, heart, and nervous system), resulting in patients experiencing cardiac, neurological, and arthritic complications (Asch et al., 1994; Singh and Girschick, 2004). A human vaccine against Lyme disease was approved by the FDA but is not currently available (Steere et al., 1998). Targeting tick proteins has the potential to disrupt tick feeding and alter *B. burgdorferi* colonization or transmission (Kurokawa et al., 2020), thereby offering a new way to interfere with the life cycle of the Lyme disease spirochete.

In the present study, we demonstrate that an *I. scapularis* adiponectin receptor-like (ISARL) protein facilitates *B. burgdorferi* colonization of the tick. ISARL-mediated stimulation of *I. scapularis* metabolic pathways are associated with spirochete colonization, and a tick complement C1q-like protein 3 contributes to ISARL activation.

## Results

### Identification and characterization of an *I. scapularis* adiponectin receptor-like protein

As tick metabolism changes during pathogen colonization, and adiponectin-associated pathways mediate diverse metabolic activities, we examined the *I. scapularis* database for two of the prominent genes linked to this pathway. The available *I. scapularis* database (taxid:6945) in NCBI was searched with the genes for mammalian adiponectin and adiponectin receptors, and results with the human and mouse genes are shown. There were no tick genes with high homology to the genes for human and mouse adiponectin full-length sequences. Interestingly, there was an *I. scapularis* gene (GenBank number: XM_029975213) with substantial homology to the human and murine adiponectin receptors, which we designated *I. scapularis* adiponectin receptor-like (*ISARL*). The corresponding ISARL protein sequence (GenBank number: XP_029831073) was also identified. The full-length *ISARL* mRNA encoded a protein with 384 amino acid residues and 71% amino acid sequence similarity to both the human and mouse adiponectin receptor protein 1 and 2. It also has high similarity (87%) to homologs from insect species, including the *Drosophila melanogaster* adiponectin receptor (GenBank number: NP_732759) (Fig. S1). Structure prediction and hydrophobicity analysis indicated that ISARL has seven transmembrane (TM) domains (Fig. S2). Comparison of the amino acid sequences between vertebrate and invertebrate species revealed that the predicted transmembrane regions are highly conserved, especially in the TM3 domain (Fig. S1).

### Silencing *ISARL* reduces *B. burgdorferi* colonization by *I. scapularis* nymphs

As *I. scapularis* lack an obvious adiponectin homolog, we examined whether expression of *ISARL* could be stimulated in the feeding vector by allowing ticks to engorge on mice, including uninfected and *B. burgdorferi*-infected animals. Interestingly a blood meal containing *B. burgdorferi* resulted in significantly increased expression of *ISARL* in the nymphal tick guts (*P* < 0.0001) (Fig. 1A). This suggests that the presence of *B. burgdorferi* in the blood meal helps to stimulate tick metabolic activity and/or that ISARL may have an important role during *B. burgdorferi* colonization of the tick gut.

**Figure 1.**
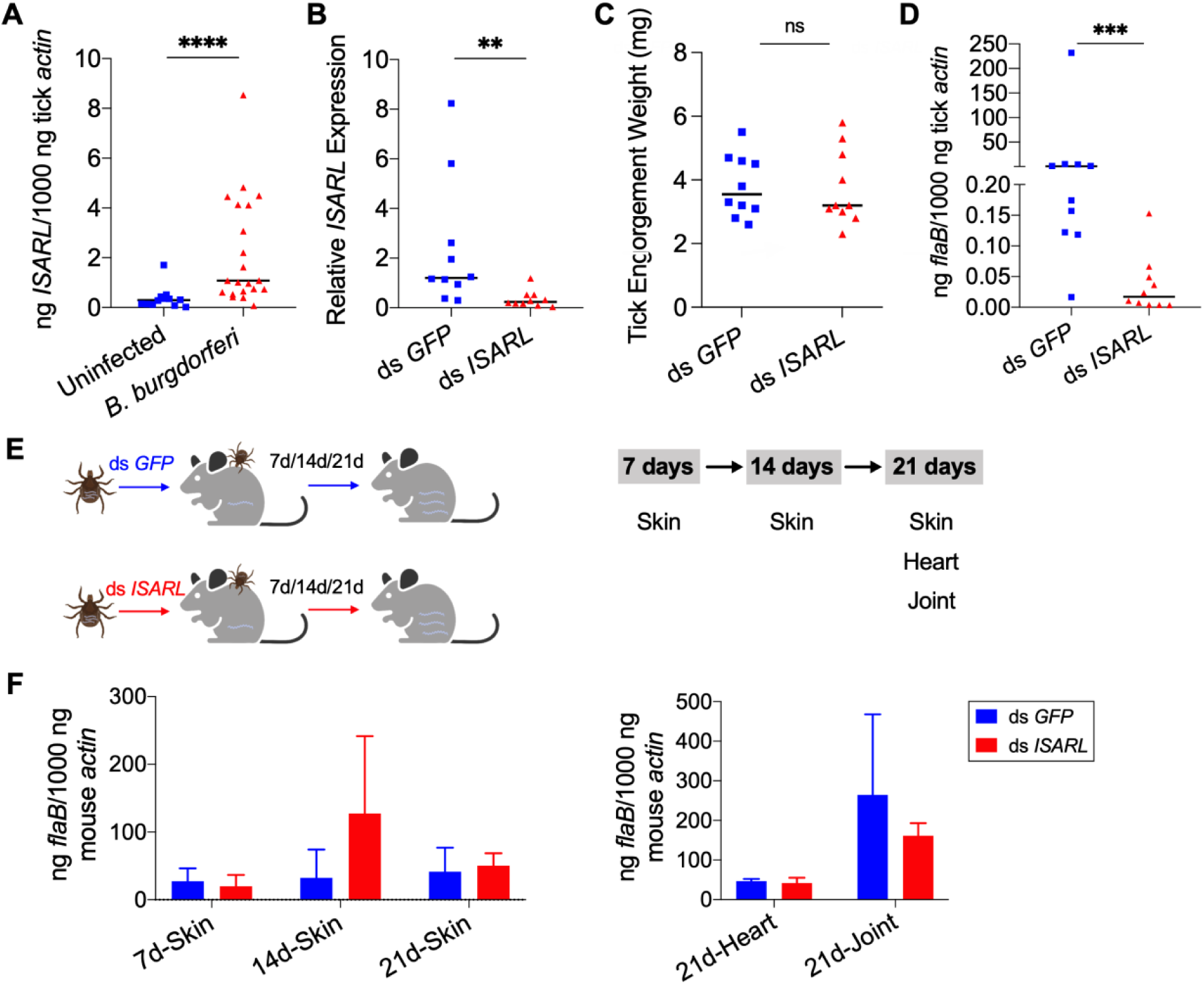
Silencing of *ISARL* significantly reduces the *B. burgdorferi* burden in nymphal tick guts. (A) *ISARL* is significantly induced in nymphal tick guts after feeding on *B. burgdorferi*-infected mice. (B) qPCR assessment of *ISARL* transcript levels following RNAi silencing of *ISARL* after feeding on *B. burgdorferi*-infected mice. (C) Nymphal engorgement weights in *ISARL*-silenced and mock-injected nymphs. Each data point represents one engorged tick. (D) qPCR assessment of *B. burgdorferi flaB* levels in guts following RNAi silencing of *ISARL* after feeding on *B. burgdorferi*-infected mice. Each data point represents one nymph gut. Horizontal bars in the above figures represent the median. Statistical significance was assessed using a non-parametric Mann-Whitney test (ns, *P* > 0.05; **, *P* < 0.01; ***, *P* < 0.001; ****, *P* < 0.0001). (E) *Borrelia*-infected nymphs microinjected with ds *ISARL* or ds *GFP* were fed on clean mice to assess transmission of the spirochete. The infection of *Borrelia* in Murine skin 7,14, and 21 days after infection, and in heart and joint tissues at 21 days was determined. (F) Murine skin 7,14, and 21 days after infection, and in heart and joint tissues at 21 days was determined by qPCR of *flaB* and normalized to mouse *actin*. Data represent the means ± standard deviations from five replicates.

Since *ISARL* expression was upregulated upon *B. burgdorferi* infection, we hypothesized that RNAi-mediated silencing of *ISARL* would affect *B. burgdorferi* colonization by nymphal *I. scapularis*. To this end, *ISARL* or *GFP* (control) dsRNA was injected into the guts of pathogen-free nymphs by anal pore injection. Then, the ticks were allowed to feed on *B. burgdorferi*-infected mice. Quantitative RT-PCR (qPCR) analysis showed a significant decrease of *ISARL* expression in the guts of ds *ISARL*-injected ticks (*P* < 0.01) when compared to that in control ds *GFP*-injected tick guts (Fig. 1B), indicating that the knockdown was successful. The engorgement weights of ds *ISARL*-injected nymphs and control ds *GFP*-injected nymphs were comparable (*P* > 0.05) (Fig. 1C), suggesting that silencing *ISARL* had no effect on tick feeding behavior. However, *ISARL*-silenced nymph guts showed a marked reduction of the *B. burgdorferi* burden (*P* < 0.001) when compared to that in control ticks (Fig. 1D), demonstrating that ISARL is associated with *B. burgdorferi* colonization in the nymphal tick gut.

### Silencing *ISARL* does not affect *B. burgdorferi* transmission by *I. scapularis* nymphs

To determine whether ISARL might also play a role in the transmission of *B. burgdorferi* to the mammalian host, we silenced *ISARL* in *B. burgdorferi*-infected nymphs by microinjection of ds *ISARL* into the ticks. The results showed that *B. burgdorferi* burdens in the skin of mice (ear skin distal from the tick bite site) at 7, 14, and 21 days post tick detachment, and in heart and joint tissues 21 days post tick detachment were comparable (*P* > 0.05) in mice fed upon by ds *GFP*- or by ds *ISARL*-injected nymphs (Figs. 1E and 1F), suggesting that silencing *ISARL* had no observable effect on *B. burgdorferi* transmission by *I. scapularis* nymphs.

### Potential ISARL-dependent pathways associated with *B. burgdorferi* colonization

To investigate the mechanisms underlying the association of ISARL with *B. burgdorferi* colonization by *I. scapularis*, we assessed the presence or absence of ISARL on tick physiology by comparing transcriptomes of ds *ISARL* and ds *GFP* (control) injected ticks after engorgement on *B. burgdorferi*-infected or uninfected mice, using RNA-seq.

After feeding on uninfected mice, 18 genes were significantly differentially expressed with six upregulated and 12 downregulated genes in the guts of ds *ISARL*- injected nymphal ticks when compared to that in control ds *GFP*-injected tick guts (Fig. 2A; Table S1). 35 genes were differentially expressed at a significant level, and all these genes were downregulated in the guts of ds *ISARL*-injected *I. scapularis* when compared to that in control ds *GFP*-injected ticks after feeding on *B. burgdorferi*-infected mice (Fig. 2B; Table S2). In addition, the transcriptome analysis further demonstrated that the *ISARL* gene was successfully silenced by RNAi (Tables S1 and S2) and this was also confirmed by qPCR (Fig. 2C). No common differentially expressed genes except *ISARL* were observed between control or experimental ticks feeding on uninfected or *B. burgdorferi*- infected mice (Fig. 2D), suggesting that the 34 significantly expressed genes were all induced by *B. burgdorferi,* or the influence of *B. burgdorferi* on the host blood components, rather than blood meal itself, in absence of ISARL.

**Figure 2.**
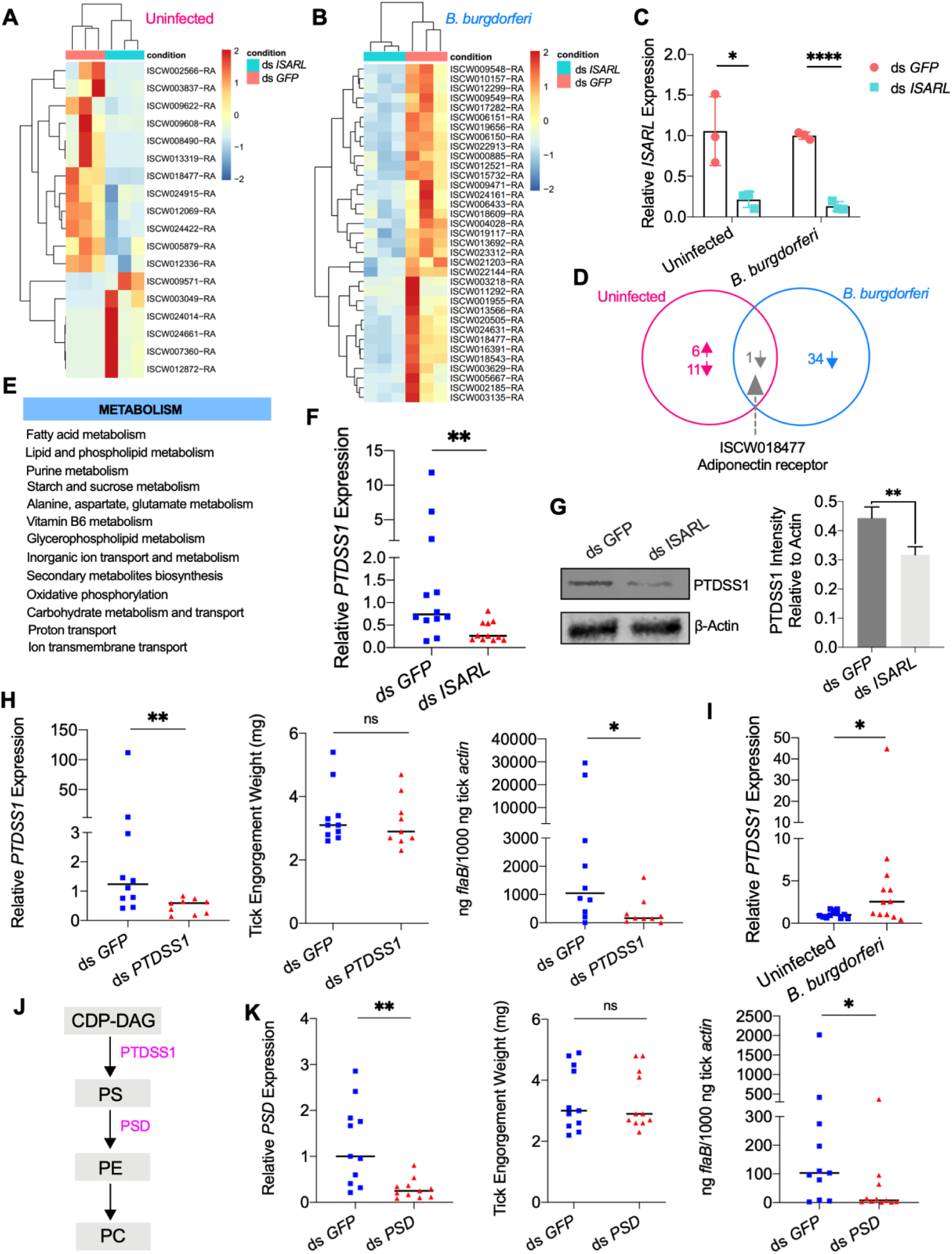
RNA-seq, qPCR validation, and RNAi-silencing assays revealed that phosphatidylserine synthase 1 (*PTDSS1*) is regulated by ISARL and is involved in *B. burgdorferi* colonization. (A) Hierarchical clustering of differentially expressed genes was generated after feeding on clean mice. (B) Hierarchical clustering of differentially expressed genes was generated after feeding on *B. burgdorferi*-infected mice. The expression levels were visualized and the scale from least abundant to highest range is from -2.0 to 2.0. The phylogenetic relationships of differentially expressed genes are shown on the left tree. The top tree indicated the cluster relationship of the sequenced samples. (C) qPCR validation of *ISARL* knockdown in tick gut. Statistical significance was assessed using a Student’s *t* test (*, *P* < 0.05; ****, *P* < 0.0001). (D) Venn diagram depicting unique and common differentially expressed genes between clean and *B. burgdorferi*-infected mice feeding. The up arrow indicated upregulation and the down arrow indicated downregulation of differentially expressed genes. (E) Metabolism pathways inferred by GO and KEGG enrichment analyses of transcriptomes comparison between ds *GFP* and ds *ISARL* injection after feeding on *B. burgdorferi*-infected mice to repletion. (F) QPCR validation of *PTDSS1* showed that *PTDSS1* is positively regulated by ISARL. (G) Western blot of PTDSS1 protein showed that PTDSS1 is positively regulated by ISARL (**, *P* < 0.05). (H) qPCR assessment of *PTDSS1* transcript level, nymphal engorgement weights, and *B. burgdorferi flaB* levels in guts following RNAi silencing of *PTDSS1* after feeding on *B. burgdorferi*-infected mice. Each data point represents one nymph. (I) *PTDSS1* is significantly induced in the nymphal tick gut after feeding on *B. burgdorferi*-infected mice. (J) PTDSS1 in involved in phospholipid pathway. Cytidine diphosphate diacylglycerol (CDP-DAG) is converted to phosphatidylserine (PS) by PTDSS1. PE: Phosphatidylethanolamine; PC: Phosphatidylcholine. (K) qPCR assessment of phosphatidylserine decarboxylase (*PSD*) transcript level, nymphal engorgement weights, and qPCR assessment of *B. burgdorferi flaB* levels in guts following RNAi silencing of *PSD* after feeding on *B. burgdorferi*-infected mice. Each data point represents one nymph. Horizontal bars in the above figures represent the median. Statistical significance was assessed using a non-parametric Mann-Whitney test (ns, *P* > 0.05; *, *P* < 0.05; **, *P* < 0.01). Source data 1. Source data for PTDSS1 protein relative quantification. Source data 2. Source data for PTDSS1 protein relative quantification.

In response to the blood meal, a significant change of the metabolic pathways in ticks was observed in absence of ISARL. In particular, based on GO functional classification and KEGG pathways analyses, the glutathione metabolic process with nine genes (e.g., gamma glutamyl transpeptidase, G2/mitotic-specific cyclin A, and glutathione S-transferase) was significantly altered. Moreover, the genes involved in metabolic pathways such propanoate metabolism and carbohydrate transport and metabolism (e.g., acyl-CoA synthetase and soluble maltase-glucoamylase) were also significantly changed in absence of ISARL after engorging on uninfected mice.

Similarly, many metabolism-associated genes, including multiple amino acids, lipids or sugars synthesis and transport genes (e.g., 3-hydroxyacyl-CoA dehydrogenase, glycogen phosphorylase, and sugar transporter) were significantly downregulated in the absence of ISARL after engorging on *B. burgdorferi*-infected mice. GO functional classification and KEGG pathways also showed that the most downregulated genes were involved in fatty acid, lipid and phospholipid, purine, amino acid, glycerophospholipid, and carbohydrate metabolism and transport pathways after silencing *ISARL* (Fig. 2E), suggesting that ISARL functions as a metabolic moderator in ticks.

### ISARL-mediated phospholipid metabolic pathways affect *B. burgdorferi* colonization

To further investigate the exact metabolism pathway(s) involving in *B. burgdorferi* colonization, we first selected 18 well-annotated and metabolism-related differentially expressed genes to validate the accuracy and reproducibility of the transcriptome bioinformatic analyses by qPCR. The samples for qPCR validation are independent of the sequencing samples. In general, the qPCR results indicated that all the tested genes showed concordant direction of change with the RNA-seq bioinformatic data except one gene, pyridoxine kinase (*PDXK*) (Fig. S3), indicating the accuracy and reliability of our RNA-seq libraries. Of these 17 down-regulated genes, four genes showed significant downregulation profiles (*P* < 0.05). These four genes included phosphatidylserine synthase I (*PTDSS1*) (Figs 2F and 2G), N-CAM Ig domain-containing protein (*NCAM*), vacuolar H+-ATPase V1 sector, subunit G (*V-ATPase*), and sideroflexin 1,2,3, putative (*SFXN*) (Fig. S4A).

Then, we silenced these four genes individually and investigated their potential roles in *B. burgdorferi* colonization. We also silenced another four genes, whose *P*-values were very close to significant (Fig. S4B). These four genes included 3-hydroxyacyl-CoA dehydrogenase, putative (*3HADH*), adenylosuccinate synthetase (*ADSS*), GMP synthase, putative (*GMPS*), and alpha-actinin, putative (*ACTN*). We did not observe a significant decrease of *B. burgdorferi* burden in nymphal tick guts after silencing *NCAM*, *V-ATPase*, *SFXN*, *ADSS*, *GMPS*, and *ACTN* compared to ds *GFP*-injected ticks (Fig. S5). Instead, we found that *PTDSS1*-silenced nymphs showed a marked reduction in the *B. burgdorferi* burden in the guts when compared to that in control ticks (*P* < 0.05) (Fig. 2H). Furthermore, a blood meal containing *B. burgdorferi* resulted in significantly increased expression of *PTDSS1* in the nymphal tick guts (*P* < 0.05) (Fig. 2I), suggesting that PTDSS1 indeed has a critical role during *B. burgdorferi* colonization of the tick gut. PTDSS1 is involved in phospholipid metabolism, and mainly uses L-serine as the phosphatidyl acceptor to generate the anionic lipid phosphatidylserine (PS), which serves as a precursor for phosphatidylethanolamine (PE) and phosphatidylcholine (PC) synthesis (Fig. 2J) (Aktas et al., 2014). Importantly, PC is one of the main phospholipids on the cellular membrane of *B. burgdorferi* (Kerstholt et al., 2020). However, *B. burgdorferi* lacks the central phospholipid metabolic enzymes. To further validate that the phospholipid metabolic pathway in tick is critical for *B. burgdorferi*, we silenced another enzyme (ISARL- unrelated), phosphatidylserine decarboxylase (*PSD*, ISCI003338), which is an important enzyme in the synthesis of PE in both prokaryotes and eukaryotes (Voelker, 1997). Interestingly, we also found a significantly decreased *B. burgdorferi* burden in ds *PSD*- injected tick guts (*P* < 0.05), and PSD and PTDSS1 elicit a similar degree of reduced *B. burgdorferi* levels (Fig. 2K). Taken together, ISARL-mediated phospholipid metabolic pathways associated with PTDSS1 have a critical role in *B. burgdorferi* colonization.

### Mammalian adiponectin regulates tick glucose metabolism pathway but has no effect on *B. burgdorferi* colonization

We further explored how the ISARL signaling pathway is activated in ticks. Although the *I. scapularis* genome encodes an adiponectin receptor homolog, an adiponectin ligand is not present, at least in currently annotated *Ixodes* genome databases. This suggests that ticks may utilize vertebrate adiponectin to activate the adiponectin receptor during a blood meal, that tick have another ligand that stimulates the receptor, or both. Since ticks are habitually exposed to adiponectin present during a bloodmeal, we examined whether the tick adiponectin receptor could interact with incoming mammalian adiponectin during blood feeding. We injected recombinant mouse adiponectin into unfed ticks and investigated whether mammalian adiponectin could activate downstream signaling of tick adiponectin receptor by RNA-seq (Fig. 3A). The data showed that one classic downstream gene of mammalian adiponectin signaling, tick glucose-6-phosphatase (*G6P*, ISCW017459), was significantly downregulated in the presence of mammalian adiponectin (Fig. 3A; Table S3). It has been demonstrated that in mammals, the binding of adiponectin to its receptor suppresses *G6P* and phosphoenolpyruvate carboxykinase (*PEPCK*) expression through an AMP-activated protein kinase (AMPK)-dependent mechanism, which further inhibits glycogenolysis and gluconeogenesis (Fig. 3B) (Tishinsky, 2012). We further searched *G6P* and *PEPCK* homologs in *I. scapularis* genome, and two *G6P* homologs (ISCW017459 and ISCW018612) and three *PEPCK* homologs (ISCW001902, and ISCW000524, ISCW001905) were identified. We designated them as *G6P1*, *G6P2, PEPCK1*, *PEPCK2*, and *PEPCK3*, respectively. We evaluated gene expression of all these five genes after injection of recombinant adiponectin and GFP proteins. Interestingly, *G6P1*, *G6P2, PEPCK2*, and *PEPCK3* were significantly downregulated in the tick gut in the presence of adiponectin (Fig. 3C). To further validate the effects on tick glucose metabolism of interaction of mammalian adiponectin and tick ISARL, we fed ticks on C57BL/6J mice deficient in adiponectin (Adipo^-/-^) and wild-type (WT) animals, and allowed them to feed to repletion (Fig. 3D). We then evaluated the expression of five *G6P* and *PEPCK* genes, and found that *G6P1* and *G6P2* also showed significant downregulation in the presence of adiponectin (*P* < 0.05), while *PEPCK* gene expression was not altered (*P* > 0.05) (Fig. 3E).

**Figure 3.**
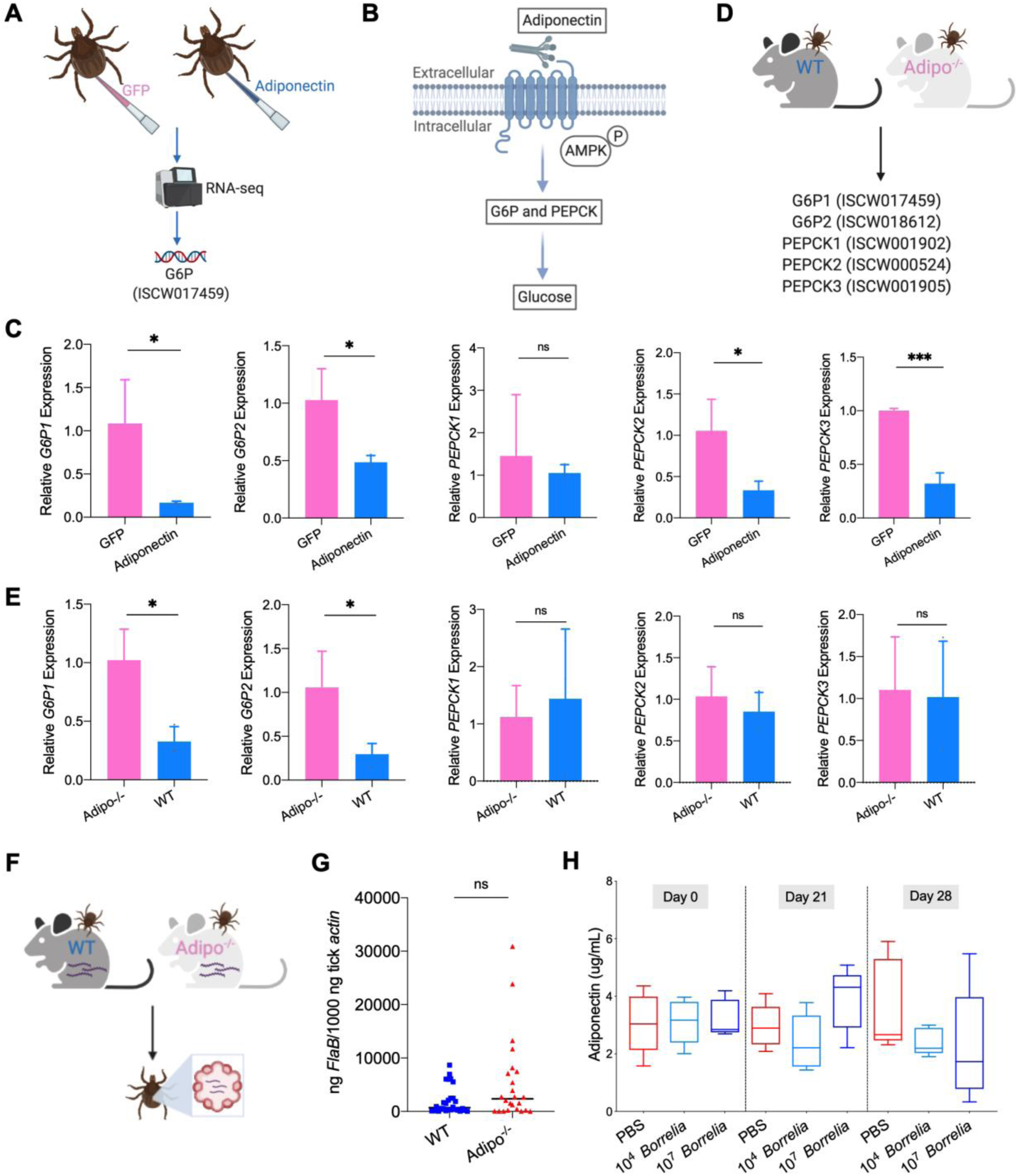
Mammalian adiponectin regulates tick glucose metabolism. (A) RNA-seq of injection of recombinant mouse adiponectin and GFP (control) proteins. One classic downstream gene of mammalian adiponectin receptor signaling, glucose-6-phosphatase (*G6P*), was significantly downregulated in the presence of mammalian adiponectin. (B) Interaction of mammal adiponectin and adiponectin receptor suppresses *G6P* and phosphoenolpyruvate carboxykinase (*PEPCK*) expression through an AMP-activated protein kinase (AMPK)-dependent mechanism, which further inhibits glycogenolysis and gluconeogenesis. (C) Injection of recombinant mouse adiponectin significantly downregulate the expression of *G6P1*, *G6P2, PEPCK2*, and *PEPCK3* in the tick gut. (D) Feed ticks on C57BL/6J WT and Adipo^-/-^ mice and then evaluate the expression of *G6P1*, *G6P2, PEPCK1*, *PEPCK2*, and *PEPCK3*. (E) After feeding on WT and Adipo^-/-^ mice, *G6P1* and *G6P2* showed significant downregulation profile in the presence of adiponectin, while *PEPCK* genes did not exhibit marked downregulation. (F) Ticks were fed on *B. burgdorferi*-infected WT and Adipo^-/-^ mice, and then *B. burgdorferi flaB* levels in guts were assessed. (G) qPCR assessment of *B. burgdorferi* burden after feeding on *B. burgdorferi*- infected WT and Adipo^-/-^ mice. No significant difference of *B. burgdorferi* burden in tick gut was observed between feeding on WT and Adipo^-/-^ mice. (H) Adiponectin concentration in mice sera following 21 and 28 days after injection of *B. burgdorferi* at the density of 10^4^ and 10^7^ cells/mL, respectively. Statistical significance was assessed using a non-parametric Mann-Whitney test (ns, *P* > 0.05; *, *P* < 0.05; **, *P* < 0.01; ***, *P* < 0.001).

To investigate whether the interaction of adiponectin and the receptor in ticks influences *B. burgdorferi* colonization, pathogen-free nymphs were placed on *B. burgdorferi*-infected WT and Adipo^-/-^ mice and allowed to feed to repletion (Fig. 3F). No significant difference of the *B. burgdorferi* burden in ticks feeding on WT and Adipo^-/-^ mice was noted (*P* > 0.05) (Fig. 3G). We also silenced the *G6P1* and *G6P2* genes to determine whether G6P-mediated glucose metabolic changes affect *B. burgdorferi* colonization. Consistent with the previous observation, there was no significant difference in the *B. burgdorferi* burden between control and *G6P1*-silenced ticks (*P* > 0.05) (Fig. S6A). *G6P2*- silenced ticks also did not show altered *B. burgdorferi* levels (*P* > 0.05) (Fig. S6B). Furthermore, the expression of *G6P1* and *G6P2* in the nymphs was not influenced by *B. burgdorferi* infection (*P* > 0.05) (Fig. S6C), suggesting that G6P1- or G6P2-mediated changes do not affect *B. burgdorferi* colonization of the tick gut. To assess any changes in the adiponectin concentration in murine serum after *B. burgdorferi* infection, the mice were injected subcutaneously with 100 uL containing 1×10^4^ or 1×10^7^ *B. burgdorferi,* or PBS as a control. We found that *B. burgdorferi* does not influence the adiponectin concentration in murine blood (Fig. 3H). Taken together, these data suggest that mammalian adiponectin can regulate ISARL-mediated glucose metabolism pathway, however, it has no effect on *B. burgdorferi* colonization.

### *B. burgdorferi* interacts with ISARL through tick C1QL3 protein

We therefore examined whether *I. scapularis* protein(s) might interact with ISARL and if *B. burgdorferi* could influence this process -- for *ISARL* silencing diminished *B. burgdorferi* colonization. To this end, we performed a blastp search of the *I. scapularis* genome with the globular C1Q domain of human and mouse adiponectin, which is known to stimulate the adiponectin receptor (Yamauchi et al., 2002). Two tick proteins had blastp hits with the human adiponectin C1Q domain (Fig. 4A) and were annotated as complement C1q-like protein 3 (C1QL3) (GenBank number: XP_002415101) and conserved hypothetical protein (GenBank number: EEC18766), respectively. These are identical proteins except that C1QL3 has a signal peptide sequence and we therefore focused on C1QL3.

**Figure 4.**
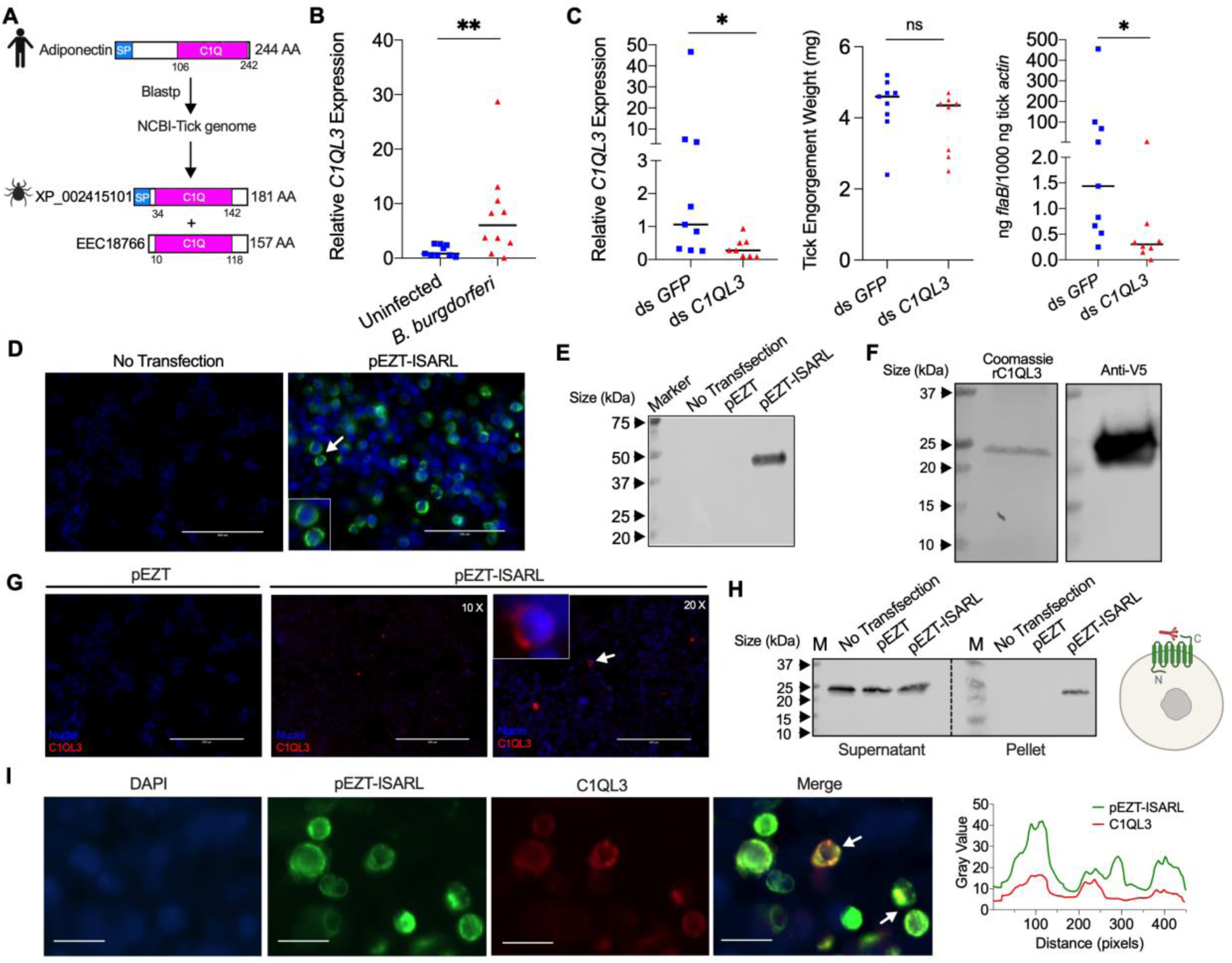
Tick complement C1q-like protein 3 (C1QL3) is involved in ISARL signaling pathways. (A) Blastp of the tick genome with the human adiponectin C1Q domain in NCBI generated two homologs and were annotated as complement C1q-like protein 3 (C1QL3) (GenBank number: XP_002415101) and conserved hypothetical protein (GenBank number: EEC18766), respectively. These are identical proteins except that C1QL3 has a signal peptide sequence. (B) *C1QL3* is significantly induced in replete nymphal tick guts after feeding on *B. burgdorferi*-infected mice. (C) qPCR assessment of *C1QL3* transcript levels, nymphal engorgement weights, and *B. burgdorferi flaB* levels in guts following RNAi silencing of *C1QL3* after feeding on *B. burgdorferi*-infected mice. (D) Human HEK293T cells were transfected with HA-tagged ISARL-expressing vector (pEZT-ISARL-HA). Forty hours post transfection, the cells were examined. The results showed that ISARL can be successfully expressed on the HEK293T cell membrane. The white arrow indicates examples of membrane expression. (E) Western blot confirmed ISARL expression in the HEK293T cells. (F) Generation of tick C1QL3 protein with His/V5-tag in a *Drosophila* expression system. Recombinant protein was assessed by SDS-PAGE gel and western blot. (G) C1QL3 is bound on the membrane of ISARL- expressed HEK293T cells. 10 X and 20 X are the microscope magnifications. The white arrow indicates one example of binding. (H) Binding of C1QL3 to ISARL as analyzed by a pull-down assay. HRP V5-tag monoclonal antibody was used to detect protein. C1QL3 was only detected in ISARL expressed cells pellet. (I) Co-immunolocalization of ISARL (green) and C1QL3 (red). The specific signal of C1QL3 protein was observed on the surface of some of ISARL expressed cells, and no signal was shown on non-successfully expressed cells membrane. The white arrows indicate examples of binding. Bar: 20 μm. The plot profile of co-localization was conducted by Image J software. Source data 1. Source data for ISARL expression. Source data 2. Source data for C1QL3 protein purification. Source data 3. Source data for C1QL3 protein purification. Source data 4. Source data for binding of C1QL3 to ISARL.

We first examined whether expression of *C1QL3* could be stimulated by *B. burgdorferi* infection. A blood meal containing *B. burgdorferi* resulted in significantly increased expression of *C1QL3* in the nymphal tick guts (*P* < 0.01) (Fig. 4B). We then generated *C1QL3*-silenced nymphs and found that these ticks had a marked reduction of the *B. burgdorferi* burden in the guts when compared to that in control *I. scapularis* (*P* < 0.05) (Fig. 4C). This is the same observation as with silencing of *ISARL*, suggesting that *B. burgdorferi* activates the ISARL-signaling pathway through the tick C1QL3 protein. Because C1QL3 C1Q domain has high similarity (64.0%) with the human adiponectin C1Q domain (Fig. S7), and C1Q proteins have been demonstrated to activate diverse pathways through the adiponectin receptor (Zheng et al., 2011), we investigated whether tick C1QL3 could interact with ISARL. Human embryonic kidney HEK293T cells were transfected with the ISARL expression vector (pEZT-ISARL). The results showed that tick ISARL can be successfully expressed, as validated by cell staining and western blot (Figs 4D and 4E), on the HEK293T cell membrane (Fig. 4D). We then generated tick C1QL3 protein in a *Drosophila* expression system (Fig. 4F). The HEK293T cells were then incubated with the recombinant C1QL3 protein. After washing and staining, recombinant C1QL3 could be detected on the surface of ISARL-expressed rather than empty plasmid transfected HEK293T cells (Fig. 4G). A pull-down assay also indicated that recombinant C1QL3 interacts with ISARL as demonstrated by the detection of C1QL3 only in ISARL expressed cells pellet (Fig. 4H). In addition, co-immunolocalization demonstrated that the C1QL3 protein specifically binds to the ISARL-expressed cell membrane (Fig. 4I). These results suggest that tick C1QL3 interacts with ISARL.

Since C1QL3 is a ligand of tick ISARL and also involved in *Borrelia* colonization, we further investigated whether C1QL3 has a role on the activation of ISARL by *Borrelia*. We first assessed if silencing of *C1QL3* influenced *ISARL* expression after feeding on *B. burgdorferi-*infected mice (Fig. 5A). QPCR assessment showed that the *ISARL* transcript level following RNAi silencing of *C1QL3* was significantly lower than that in control ds *GFP*-injected tick guts after feeding on *B. burgdorferi*-infected mice (*P* < 0.05) (Fig. 5B). More importantly, after silencing *C1QL3*, a blood meal containing *B. burgdorferi* did not significantly increase expression of *ISARL* in the nymphal tick guts as compared to feeding on clean mice (*P* > 0.05) (Fig. 5C), further suggesting that C1QL3 plays a role in the ISARL signaling pathways.

**Figure 5.**
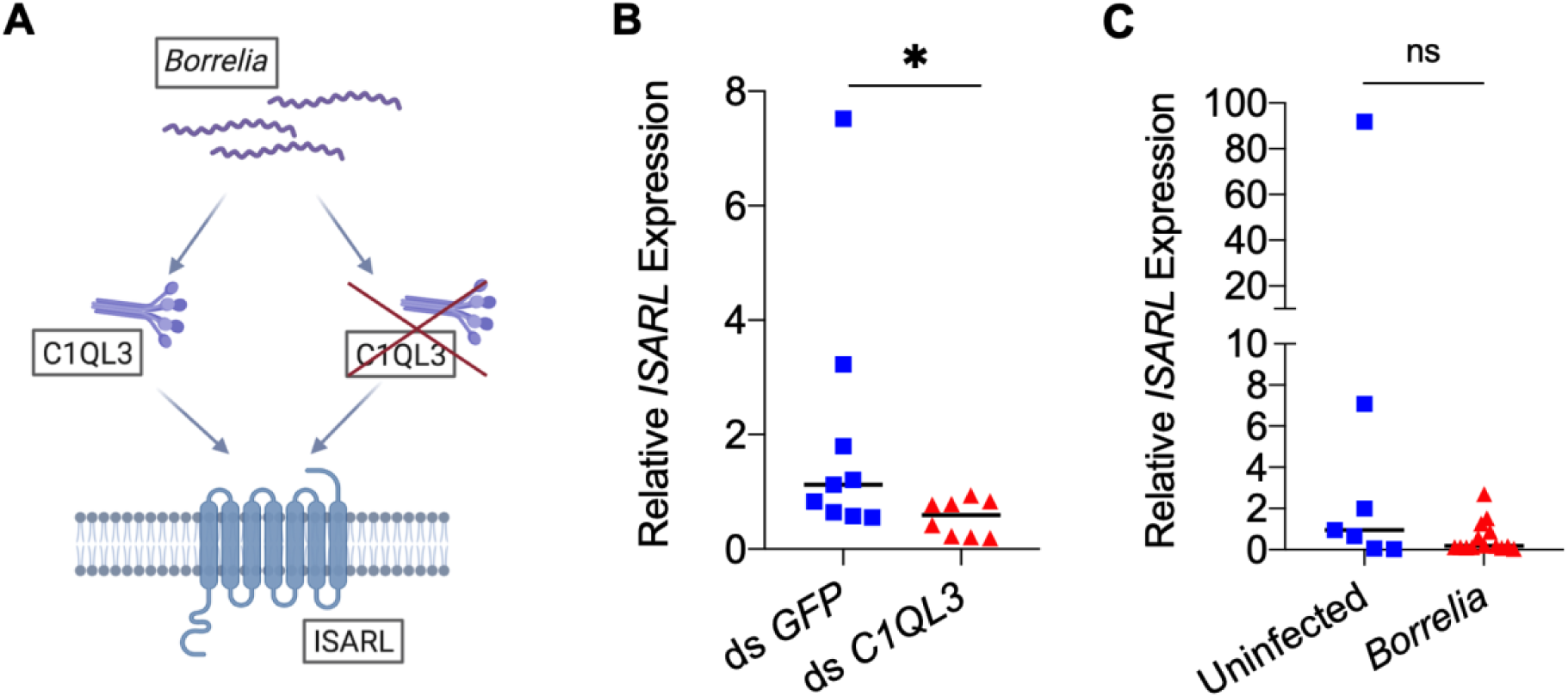
C1QL3 plays a role on the activation of ISARL by *Borrelia*. (A) Analysis of how silencing of *C1QL3* influences *ISARL* expression after feeding on *B. burgdorferi-* infected mice. (B) qPCR assessment showed that *ISARL* transcript levels following RNAi silencing of *C1QL3* were significantly lower than that in control ds *GFP*-injected tick guts after feeding on *B. burgdorferi*-infected mice. (C) qPCR assessment showed that a blood meal containing *B. burgdorferi* did not significantly increase expression of *ISARL* in the nymphal tick guts as compared to feeding on clean mice after RNAi silencing of *C1QL3*. Each data point represents one nymph. Horizontal bars in the above figures represent the median. Statistical significance was assessed using a non-parametric Mann-Whitney test (ns, *P* > 0.05; *, *P* < 0.05).

## Discussion

Adiponectin is a hormone, secreted mainly from adipocytes, that stimulates glucose utilization and fatty acid oxidation (Berg et al., 2001; Fruebis et al., 2001). The key roles of adiponectin in regulating energy homeostasis are mediated by adiponectin receptors across species including humans, yeast, nematodes, and flies (Kwak et al., 2013; Narasimhan et al., 2005; Svensson et al., 2011; Yamauchi et al., 2003). In this study, we have identified and characterized an adiponectin receptor homologue from *I. scapularis*, ISARL. ISARL shares significant sequence similarities with human, mouse, and *Drosophila* adiponectin receptors. In addition, ISARL contains the canonical features of adiponectin receptors, including conserved TM domains, a long internal N-terminal region, and a relatively short external C-terminal region. The highly conserved amino acids and the structures of ISARL and the receptor from *D. melanogaster* suggest that their ligands and signaling pathways may also be conserved in arthropods. However, homologs of adiponectin have not yet been identified in arthropods, suggesting that ligands for adiponectin receptors in arthropods may interact in different ways that in vertebrates.

The Lyme disease agent, *B. burgdorferi*, engages in intimate interactions with *I. scapularis* during its acquisition and colonization of the tick gut (Radolf et al., 2012). This is accompanied by dramatic changes in the expression profiles of *Borrelia* and tick gut genes, which are critical drivers for colonization, persistence or transmission (Kurokawa et al., 2020; Narasimhan et al., 2017). In our study, expression of *ISARL* was significantly increased in the nymphal tick gut after *B. burgdorferi* infection. The upregulation of *ISARL* correlates with *Borrelia* infection in the gut. More interestingly, after silencing *ISARL* expression in the tick gut by anal pore injection, nymphal tick guts showed a marked reduction in the *B. burgdorferi* burden when compared to that in control ticks, demonstrating that ISARL facilitates *B. burgdorferi* colonization.

We utilized RNA-seq to elucidate the pathways that are altered when *ISARL* is silenced in ticks that engorge on clean and *B. burgdorferi*-infected mice. All the differentially expressed genes were downregulated, and GO functional classification and KEGG pathways also showed that the most downregulated genes are involved in fatty acid, lipid and phospholipid, purine, amino acid, glycerophospholipid, and carbohydrate metabolism and transport pathways. Therefore, ISARL in ticks functions as a metabolic regulator.

Importantly, ISARL can regulate a critical enzyme involved in phospholipid metabolism, PTDSS1. Regulation of PTDSS1 by adiponectin receptors is also found in other organisms such as yeast, where the adiponectin receptor homolog Izh2 is connected to phospholipid metabolism through co-regulation of the expression of inositol-3-phosphate synthase (*INO1*) and phosphatidylserine synthase (*CHO1*, homolog of *PTDSS1*) genes with zinc-responsive activator protein (*Zap1*) (Ušaj et al., 2015). Silencing of *I. scapularis PTDSS1* led to a reduced spirochete burden in the guts, thereby linking *B. burgdorferi* colonization with phospholipid metabolism. The *B. burgdorferi* genome is small and encodes a limited number of metabolic enzymes (Casjens et al., 2000; Fraser et al., 1997). The restricted biosynthetic capability forces *B. burgdorferi* to rely on its vertebrate and arthropod hosts for nutrients or enzymes that it cannot synthesize (Tilly et al., 2008). Interestingly, we also found that silencing of *I. scapularis 3HADH,* which is involved in fatty acid metabolic processes, decreased the *B. burgdorferi* burden in tick gut (Figure S5D). The markedly decreased *B. burgdorferi* burden in ticks after silencing of *PTDSS1*, *PSD* and *3HADH*, suggests that the spirochete may require the tick for selected metabolic needs.

We also found that *B. burgdorferi* can interact with an adiponectin-related protein, C1QL3, in ticks, which associates with ISARL. The interactions lead to phospholipid metabolism changes in ticks. We propose that C1QL3 in tick is mainly involved in metabolism, rather than complement activation, as demonstrated by the decreased *B. burgdorferi* level after silencing C1QL3. Indeed, some of C1Q/TNF family proteins are associated with metabolism. In addition to adiponectin, proteins such as C1Q/TNF-related protein 3 (CTRP3), CTRP5, CTRP9, CTRP13 (C1QL3), and CTRP15 also belong to adipokine family, and have been reported to be associated with the regulation of glucose, lipid or other metabolisms (Jiang et al., 2018; Li et al., 2011; Mi et al., 2019; Wei et al., 2011; Wolf et al., 2016). Importantly, C1Q proteins have been demonstrated to activate diverse pathways through adiponectin receptor (Zheng et al., 2011).

In summary, we demonstrate that ISARL plays a key role in metabolic pathways in *I. scapularis*. ISARL-mediated phospholipid metabolism by PTDSS1 contributes to *B. burgdorferi* colonization and an adiponectin-related protein, C1QL3, is involved in ISARL signaling pathway. These studies elucidate a new pathway involved in tick metabolism, and demonstrate that *B. burgdorferi* co-opts the activation of this pathway to facilitate colonization of *I. scapularis*. These processes are crucial to understanding the complex life cycle of the Lyme disease agent within ticks, and may be applicable to other arthropods and arthropod-borne infectious agents.

## Materials and Methods

### Ethics statement

Animal care and housing were performed according to the Guide for the Care and Use of laboratory Animals of National Institutes of Health, USA. All protocols in this study were approved by the Yale University Institutional Animal Care and Use Committee (YUIACUC) (approval number 2018-07941). All animal experiments were performed in a Biosafety Level 2 animal facility.

### Mice, spirochetes, ticks, and cells

C3H/HeJ mice, C57BL/6J mice wild-type (WT), and C57BL/6J mice deficient in adiponectin (Adipo^-/-^) were purchased from the Jackson Laboratory (https://www.jax.org/strain/008195). All mice were bred and maintained in a pathogen-free facility at Yale University. The spirochetes *B. burgdorferi* N40 were grown at 33°C in Barbour-Stoenner-Kelly H (BSK-H) complete medium (Sigma-Aldrich, #B8291) with 6% rabbit serum. The live cell density was ∼10^6^-10^7^ cells/mL as determined by dark field microscopy and hemocytometric analysis. To obtain *B. burgdorferi*-infected mice, the mice were injected subcutaneously with 100 uL of *B. burgdorferi* N40 (1×10^5^ cells/mL). Two weeks after inoculation, the *B. burgdorferi* burden in mice was assayed by qPCR analysis of spirochete DNA in murine ear punch biopsies as described below. DNA was extracted from mouse skin-punch biopsies using the DNeasy tissue kit (QIAGEN, #69506) according to the manufacturer’s protocol. The DNA was analyzed by qPCR using *flagellinB* (*flaB*) primers, and data was normalized to mouse *actin*. The primer sequences are shown in Table S4. Pathogen-free *I. scapularis* larvae were acquired from the Centers for Disease Control and Prevention. The larval ticks were fed to repletion on pathogen-free C3H/HeJ mice and allowed to molt to nymphs. *B. burgdorferi*-infected nymphs were generated by placing larvae on *B. burgdorferi*-infected C3H/HeJ mice, and fed larvae were molted to nymphs. Nymphal ticks were maintained at 85% relative humidity with a 14h light and 10h dark period at 23 °C. Human embryonic kidney HEK293T cells (ATCC, #CRL-3216) was used for vitro studies. The HEK293T cells were grown in Dulbecco’s Modified Eagle’s Medium (DMEM, ThermoFisher, #11965-118) media supplemented with 10% Fetal Bovine Serum (FBS, Sigma, #12306C-500).

### Identification and characterization of the *I. scapularis* adiponectin receptor-like (*ISARL*) gene

The human adiponectin receptor protein 1 (GenBank number: NP_001277482) and 2 (GenBank number: NP_001362293) sequences were used to conduct tblastn and blastp searches against the available black-legged tick database (taxid:6945) using NCBI default parameters. Tick adiponectin receptor sequence was further validated by amplification with primers in Table S4. Multiple alignment of protein sequences were performed using the Clustal Omega (https://www.ebi.ac.uk/Tools/msa/clustalo/) (Madeira et al., 2019) or Uniprot (https://www.uniprot.org/align/). The similarities of adiponectin receptor protein sequences were measured in EMBOSS supermatcher (https://www.bioinformatics.nl/cgi-bin/emboss/supermatcher). The protein structure of ISARL was predicted in SWISS-MODEL (https://swissmodel.expasy.org/) (Guex and Peitsch, 1997; Waterhouse et al., 2018). Hydrophobicity analysis was performed using ProtScale (https://web.expasy.org/protscale/) (Gasteiger et al., 2005).

### Tick exposure to *B. burgdorferi* and expression of *ISARL*

To evaluate gene expression of *ISARL* upon *B. burgdorferi* infection, pathogen-free *I. scapularis* nymphs were placed on *B. burgdorferi-*free and -infected mice (C3H/HeJ). At least three mice were used in each experiment, and the ticks were allowed to feed to repletion. Both *B. burgdorferi*-free and -exposed tick guts were dissected under the dissecting microscope. The RNA from dissected guts was purified by Trizol (Invitrogen, #15596-018) following the manufacturer’s protocol, and cDNA was synthesized using the iScript cDNA Synthesis Kits (Bio-Rad, #1708891). qPCR was performed using iQ SYBR Green Supermix (Bio-Rad, #1725124) on a Bio-Rad cycler with a program consisting of an initial denaturing step of 2 min at 95°C and 45 amplification cycles consisting of 20 s at 95°C followed by 15 s at 60°C, and 30 s at 72°C. The genes and corresponding primer sequences are shown in Table S4. The specific target transcripts of *ISARL* and the reference gene tick *actin* were quantified by extrapolation from a standard curve derived from a series of known DNA dilutions of each target gene, and data was normalized to tick *actin*.

### RNAi silencing of targeted genes

Fed-nymph gut cDNA was prepared as described above and used as template to amplify segments of targeted genes. The PCR primers with T7 promoter sequences are shown in Table S4. Double-stranded RNA (dsRNA) were synthesized using the MEGAscript RNAi kit (Invitrogen, #AM1626M) using PCR-generated DNA template that contained the T7 promoter sequence at both ends. The dsRNA quality was examined by agarose gel electrophoresis. DsRNA of the *Aequorea victoria* green fluorescent protein (GFP) was used as a control. Pathogen-free and -infected tick nymphs were injected in the anal pore with dsRNA (6 nL) using glass capillary needles as described by Narasimhan and colleagues (2004).

### Effects of silenced genes on *B. burgdorferi* colonization and transmission

To examine the effects of silencing targeted genes on the colonization of *B. burgdorferi* in the tick gut, dsRNA microinjected pathogen-free *I. scapularis* nymphs were placed on *B. burgdorferi*-infected mice (C3H/HeJ) and allowed to feed to repletion. The ticks were then collected for gut dissection. The *B. burgdorferi* burden in the tick gut was quantified by amplifying *flaB*. *FlaB* was quantified by extrapolation from a standard curve derived from a series of known DNA dilutions of *flaB* gene, and data was normalized to tick *actin*. The knockdown efficiency of targeted genes was tested as described above. Specifically, the expression of targeted genes was estimated with the ΔΔC_T_ method (Schmittgen and Livak, 2008) using the reference gene *actin*. To test the effects of silencing *ISARL* on the transmission of *B. burgdorferi*, a group of three to five *GFP* or *ISARL* dsRNA injected *B. burgdorferi*-infected nymphs were placed on each C3H/HeJ mouse (at least five mice each in the *GFP* or *ISARL* dsRNA groups) and allowed to feed to repletion. Ticks are placed on the mouse head/back between the ears. At 7 and 14 days-post tick detachment, the mice were anesthetized, and skin was aseptically punch biopsied and assessed for spirochete burden by qPCR. Ticks feed in head area and skin punch biopsies are collected from the pinnae /ears. This site is considered distal as it is not at the site of tick bite. Twenty-one days post tick detachment, the mice were sacrificed, and ear skin, heart, and joints were aseptically collected and assessed for spirochete burden by qPCR.

### RNA-seq and bioinformatic analyses

DsRNA (ds *ISARL* and ds *GFP*) microinjected pathogen-free *I. scapularis* nymphs were placed on clean and *B. burgdorferi*-infected mice (C3H/HeJ), respectively, and allowed to feed to repletion. Then, the ticks were collected for gut dissection. Total RNA was purified as described above. In addition, to check the transcriptional alterations in the tick gut in the presence of mammalian adiponectin, pathogen-free tick nymphs were injected in the anal pore with recombinant mouse adiponectin (MilliporeSigma, #SRP3297) and GFP proteins (SinoBiological, #13105-S07E). Then, the guts were dissected after 8h injection, and RNA was purified. The RNA samples were then submitted for library preparation using TruSeq (Illumina, San Diego, CA, USA) and sequenced using Illumina HiSeq 2500 by paired-end sequencing at the Yale Centre for Genome Analysis (YCGA). The *I. scapularis* transcript data was downloaded from the VectorBase (https://vectorbase.org/vectorbase/app/) (Giraldo-Calderon et al., 2015) and indexed using the kallisto-index (Bray et al., 2016). The reads from the sequencer were pseudo-aligned with the index reference transcriptome using kallisto (Bray et al., 2016). The counts generated were processed by DESeq2 (Love et al., 2014) in RStudio (https://rstudio.com). Gene ontology (GO) enrichment analysis and Kyoto Encyclopedia of Genes and Genomes (KEGG) pathway enrichment analyses were conducted using the functional annotation tool DAVID 6.8 (Sherman and Lempicki, 2009).

Recombinant mouse adiponectin (Sigma, #SRP3297) and GFP protein (Sino Biological, #13105-S07E) were injected into pathogen-free *I. scapularis* nymphs. After 8h, the ticks were collected for gut dissection. Total RNA was purified and RNA-seq and analyses were performed as described above.

### Expression of ISARL and binding assays

Tick *ISARL* gene was PCR amplified from nymph cDNA using the primer pair listed in Table S4, then cloned into the *Xba*I and *Not*I sites of the pEZT-Dlux, a modified pEZT-BM vector (Addgene, #74099) in-frame with a HA-tag sequence, by Gibson Assembly Cloning Kit (NEB, #E5510S). HEK293T cells were transfected with the *ISARL* expression plasmid (pEZT-ISARL) using TransIT 2020 (Mirus, #MIR5404). After 40 h post transfection, the cells were washed with 1X PBS and then incubated with rC1QL3 protein with His/V5 tag, respectively. After 16 h incubation with gentle agitation, the cells were washed with PBS and fixed in 4% PFA for 15 min at room temperature. Then, the cells were blocked in 1% BSA in PBS for 1 h, and subsequently immunolabeled with anti-HA antibody (1:100, Cell Signaling Technology, #C29F4) for checking ISARL expression, and V5 tag monoclonal antibody (1:100, Invitrogen, # R960-25) for checking C1QL3 binding. Cells were washed with PBS three times and then immunolabeled with secondary antibodies of Goat anti-Rabbit IgG (H+L) Highly Cross-Adsorbed Secondary Antibody, Alexa Fluor 488 (1:100, Invitrogen, #A-11034) and Goat anti-Mouse IgG (H+L) Cross-Adsorbed Secondary Antibody, Alexa Fluor 555 (1:100, Invitrogen, #A-21422) for 1 h at room temperature. Nuclei were stained with DAPI (Invitrogen, #D9542). After staining, the fluorescence signals were examined with an EVOS FL Auto Cell Imaging System (Thermo Fisher Scientific). We also conducted plot profile to help analyze co-localization by Image J software.

For checking ISARL expression by western blot, after 40 h post transfection, the cells were washed with 1X PBS and then lysed with 4X Laemmli Sample Buffer (Bio-Rad, #1610747). After centrifuge at high speed, the supernatant was loaded to perform western blot as described below. HRP Anti-His tag antibody (1:10,000, abcam, #ab3553) or anti-HA antibody (1:1000, Cell Signaling Technology, #C29F4) was used to detect expression of ISARL.

We conducted a pull-down assay to check the binding of ISARL and C1QL3 as described in Schuijt et al. (2011a). Briefly, HEK293T cells were transfected as described above. After 40 h post transfection, the cells were washed and suspended with 1X PBS and then incubated with rC1QL3 protein for 16 h with gentle agitation, respectively. Then the cells were pelleted, and the pellet and supernatant were separated. The pellet was washed five to eight times in 1.5 ml PBS/0.1% BSA and was resuspended in the same volume as the supernatant. Equal volumes of supernatant and pellet were used to run western blot as described below. HRP V5-tag monoclonal antibody (1:1000, Invitrogen, # R961-25) was used to detect protein.

### Adiponectin concentration in serum after *B. burgdorferi* infection

To assess the adiponectin concentration change in mice serum after *B. burgdorferi* infection, the C3H/HeJ mice were injected subcutaneously with 100 uL 1×10^4^ and 1×10^7^ cells/mL *B. burgdorferi* and PBS as a control (five mice in each group). At 0, 21 and 28 days-post inoculation, the blood was collected from mice. The sera were separated from mice blood samples by centrifugation at 1000 x g for 10 min at 4 °C. The adiponectin in mice serum was quantified by Mouse Adiponectin/Acrp30 Quantikine ELISA Kit (R&D Systems, #MRP300).

### Effects of adiponectin in mice blood on *B. burgdorferi* colonization

Pathogen-free *I. scapularis* nymphs were placed on *B. burgdorferi*-infected WT and Adipo^-/-^ mice (C57BL/6J) and allowed to feed to repletion. The ticks were then collected for gut dissection. The *B. burgdorferi* burden in the tick gut was quantified as described above.

### Purification of recombinant proteins

The *C1QL3* was PCR amplified from tick nymph cDNA using the primer pair listed in Table S4, then cloned into the *Bgl*II and *Xho*I sites of the pMT/BiP/V5-His vector (Invitrogen, #V413020). The recombinant protein was expressed and purified using the *Drosophila* Expression System as described previously (Schuijt et al., 2011b). The protein was purified from the supernatant by TALON metal affinity resin (Clontech, #635606) and eluted with 150 mM imidazole. The eluted samples were filtered through a 0.22-μm filter and concentrated with a 10-kDa concentrator (MilliporeSigma, #Z740203) by centrifugation at 4 °C. Recombinant protein purities were assessed by SDS-PAGE using 4-20% Mini-Protean TGX gels (Bio-Rad, #4561094) and quantified using the BCA Protein Estimation kit (ThermoFisher Scientific, #23225).

### Western blot

Proteins were separated by SDS-PAGE at 160 V for 1h. Proteins were transferred onto a 0.45-m-pore-size polyvinylidene difluoride (PVDF) membrane (Bio-Rad, #1620177) and processed for immunoblotting. The blots were blocked in 1% non-fat milk in PBS for 60 min. Primary antibodies of PTDSS1 Rabbit pAb (1:1000, Abclonal, #A13065), Anti-beta Actin antibody (1:1000, abcam, #ab8224), HRP Anti-6X His tag antibody (1:10,000, abcam, #ab3553) or HRP V5 tag monoclonal antibody (1:1000, Invitrogen, # R961-25) were diluted in 0.05% PBST and incubated with the blots for 1 h at room temperature or 4 °C overnight. HRP-conjugated secondary antibody (1:2500, Invitrogen, #62-6520 and #31466) were diluted in PBST and incubated for 1 h at room temperature. After washing with PBST, the immunoblots were imaged and quantified with a LI-COR Odyssey imaging system.

### Statistical analysis

Statistical significance of differences observed in experimental and control groups was analyzed using GraphPad Prism version 8.0 (GraphPad Software, Inc., San Diego, CA). Non-parametric Mann-Whitney test or unpaired t test were utilized to compare the mean values of control and tested groups, and *P* < 0.05 was considered significant. The exact *P* values are shown in the source data.

## ACKNOWLEDGEMENTS

This work was support by grants from the NIH (AI126033, AI138949) and the Steven and Alexandra Cohen Foundation. We sincerely thank Ms Kathleen DePonte for her excellent technical assistance. We would like to acknowledge that figures were created using BioRender.

## AUTHOR CONTRIBUTIONS

X.T. and E.F. conceived and shaped the overall direction of the project. X.T. Y.C. G.A. J.H. A.S. A.M. Y.M.C. M.J.W. C.L.B. and H.M. performed experiments. Y.C. and J.H. bred the adiponectin knockout mice. S.M. conducted RNA-seq data analyses. S.N. and U.P. was involved in the critical discussion of this study. Illustrations were created by X.T. using BioRender. X.T. and E.F. wrote the manuscript with feedback and discussions from all co-authors.

## LEAD CONTACT

Further information and requests for resources and reagents should be directed to and will be fulfilled by the Lead Contact, Erol Fikrig

## DATA AVAILABILITY

The RNA-seq data are available in the Gene Expression Omnibus (GEO) repository at the National Center for Biotechnology Information under the accession number: GSE169293.

## Supplemental information

**Figure S1.**
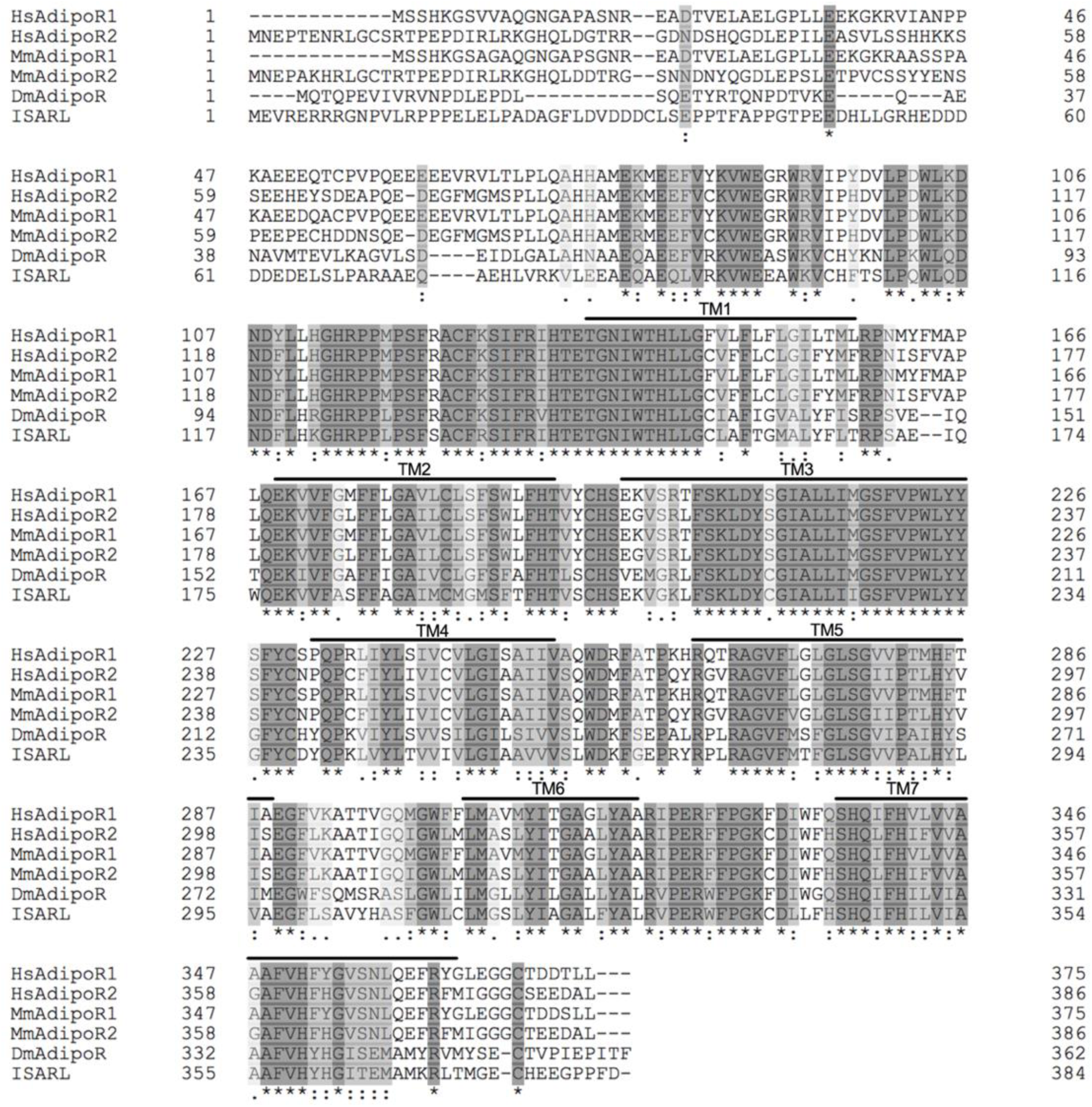
Protein sequence comparison of adiponectin receptors. Multiple sequence alignment of *Ixodes scapularis* ISARL with the amino acid sequences of homologs identified in *Homo sapiens* (NP_001277482, HsAdipoR1; NP_001362293, HsAdipoR2), *Mus musculus* (NP_001292998, MmAdipoR1; NP_001342621, MmAdipoR2), and *Drosophila melanogaster* (NP_732759, DmAdipoR). Seven transmembrane (TM1-TM7) domain regions are marked by upper lines. * indicates positions which have a single, fully conserved residue (dark grey). : indicates conservation between groups of strongly similar properties (light grey). . indicates conservation between groups of weakly similar properties (white grey). The TM domains is based on the experimentally defined human adiponectin receptors.

**Figure S2.**
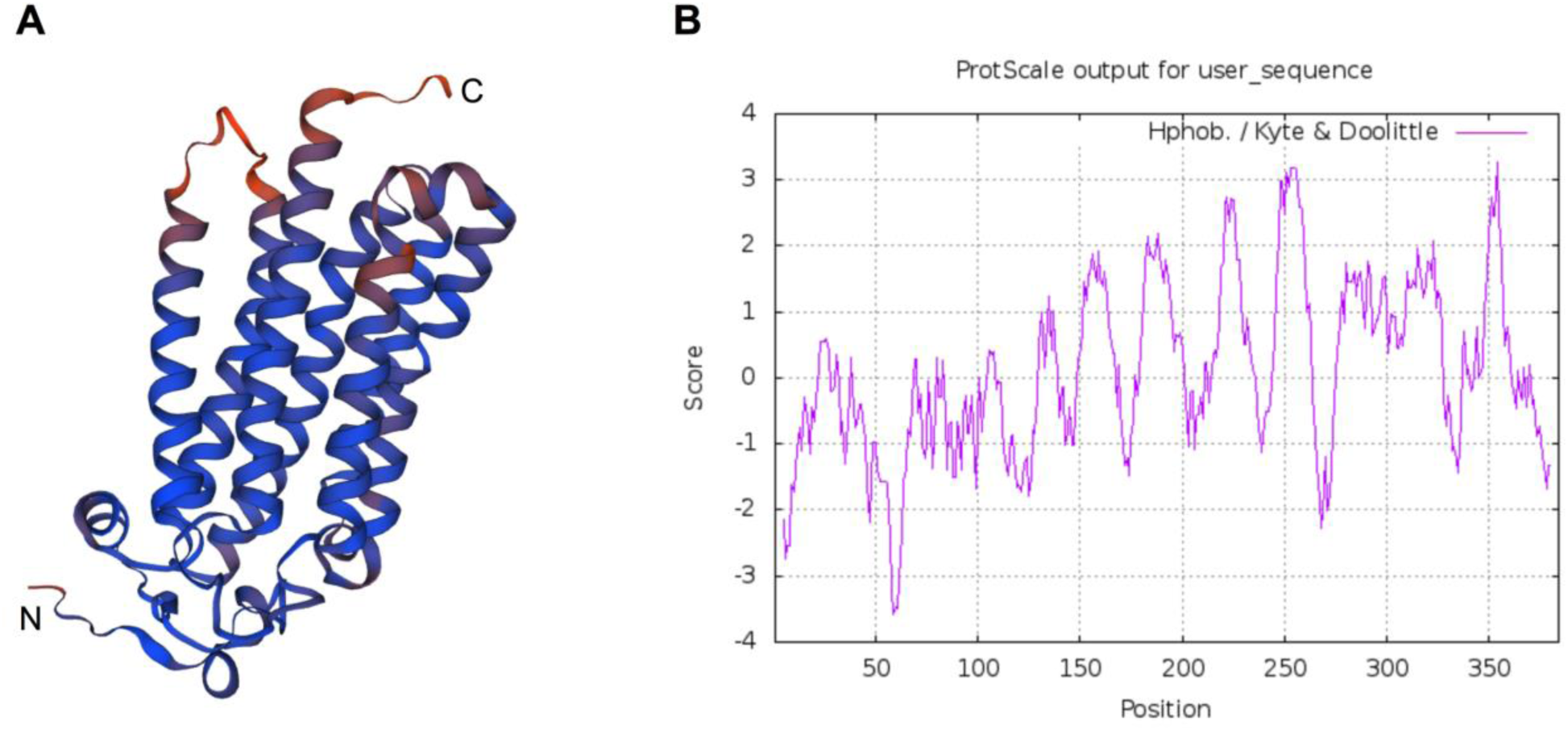
Predicted protein structure and hydrophobicity of ISARL. Seven transmembrane (TM) domains were identified in the ISARL protein based on (A) protein structure prediction and (B) hydrophobicity analysis.

**Figure S3.**
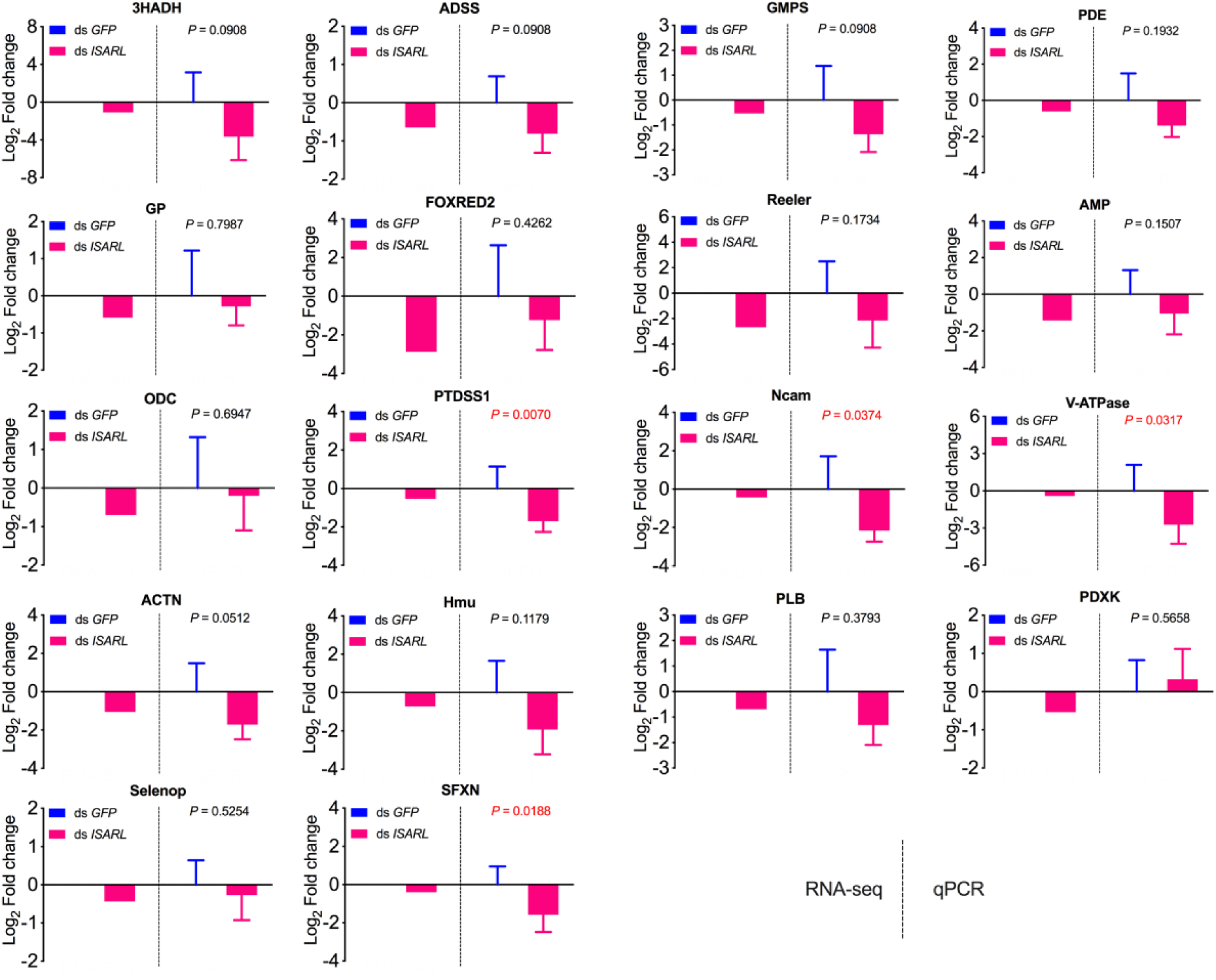
QPCR validation of 18 well-annotated and metabolism-related differentially expressed genes. 3HADH, 3-hydroxyacyl-CoA dehydrogenase, putative; ADSS, Adenylosuccinate synthetase; GMPS, GMP synthase, putative; PDE, cAMP and cAMP-inhibited cGMP 3,5-cyclic phosphodiesterase; GP, glycogen phosphorylase; FOXRED2, FAD dependent oxidoreductase domain-containing protein 2; Reeler, Secreted protein with Reeler domain; AMP, AMP dependent CoA ligase; ODC, Oxodicarboxylate carrier protein; PTDSS1, Phosphatidylserine synthase I; Ncam, N-CAM Ig domain-containing protein; V-ATPase, vacuolar H+-ATPase V1 sector, subunit G; ACTN, Alpha-actinin, putative; Hmu, Hemomucin, putative; PLB, Phospholipase B-like; PDXK, Pyridoxine kinase, putative; Selenop, selenoprotein P precursor; SFXN, sideroflexin 1,2,3, putative.

**Figure S4.**
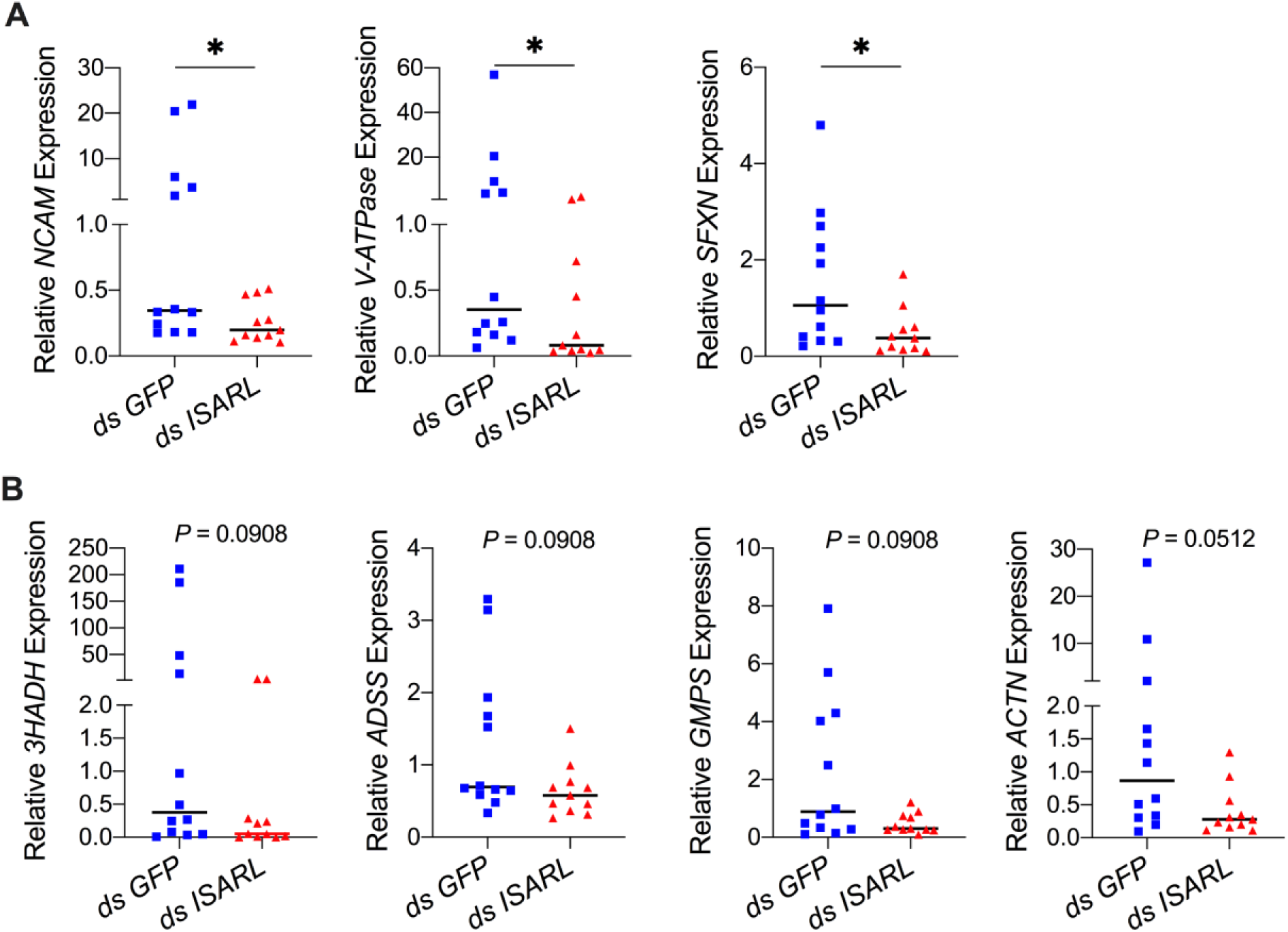
QPCR validation of differentially expressed genes from the RNA-seq dataset. (A) N-CAM Ig domain-containing protein (*NCAM*), Vacuolar H+-ATPase V1 sector, subunit G (*V-ATPase*), and Sideroflexin 1,2,3, putative (*SFXN*) were significantly downregulated following RNAi silencing of *ISARL* after feeding on *B. burgdorferi*-infected mice. (B) 3-hydroxyacyl-CoA dehydrogenase, putative (*3HADH*), Adenylosuccinate synthetase (*ADSS*), GMP synthase, putative (*GMPS*), and Alpha-actinin, putative (*ACTN*) were downregulated following RNAi silencing of *ISARL* after feeding on *B. burgdorferi*- infected mice (*P*-values are close to significant of 0.05). Each data point represents one nymph gut. Horizontal bars in the above figures represent the median. Statistical significance was assessed using a non-parametric Mann-Whitney test (*, *P* < 0.05).

**Figure S5.**
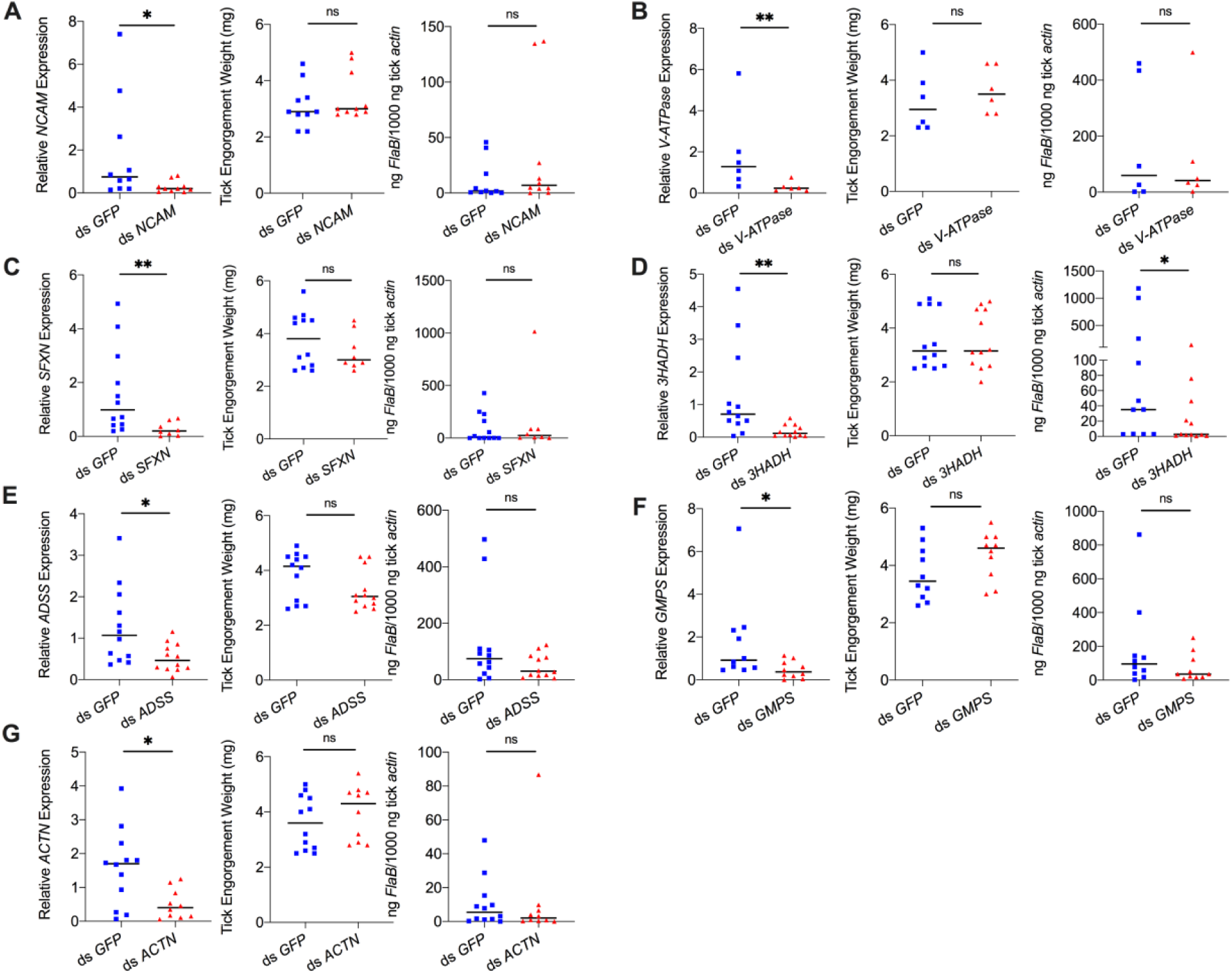
Silencing of differentially expressed genes and effects on *B. burgdorferi* acquisition. Silencing of (A) *NCAM*, (B) *V-ATPase*, (C) *SFXN*, (E) *ADSS*, (F) *GMPS*, and (G) *ACTN* has no effect on *B. burgdorferi* acquisition. Silencing of (D) *3HADH* decreased the *B. burgdorferi* burden in tick gut. 3HADH is involved in fatty acid metabolic processes, suggesting that tick fatty acid metabolism may also influence acquisition of *B. burgdorferi*. 3HADH is not significantly regulated by ISARL, it was not considered further in this study. Each data point represents one nymph gut. Horizontal bars in the above figures represent the median. Statistical significance was assessed using a non-parametric Mann-Whitney test (ns, *P* > 0.05; *, *P* < 0.05; **, *P* < 0.01).

**Figure S6.**
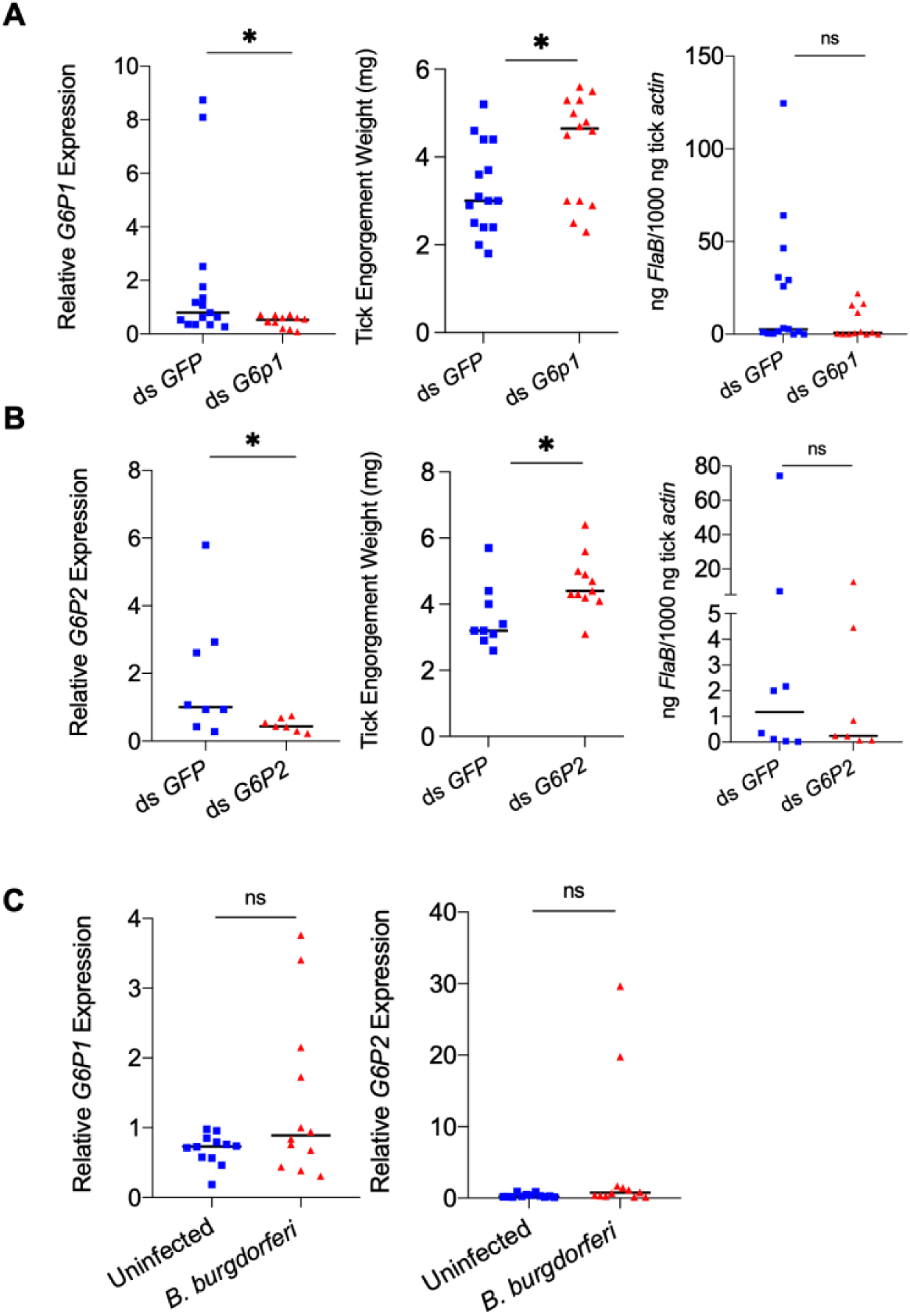
Mammalian adiponectin and tick glucose metabolism changes have no effect on *B. burgdorferi* acquisition. (A) qPCR assessment of *G6P1* transcript level, nymphal engorgement weights, and qPCR assessment of *B. burgdorferi flaB* levels in guts following RNAi silencing of *G6P1* after feeding on *B. burgdorferi*-infected mice. (B) qPCR assessment of *G6P2* transcript level, nymphal engorgement weights, and qPCR assessment of *B. burgdorferi flaB* levels in guts following RNAi silencing of *G6P2* after feeding on *B. burgdorferi*-infected mice. (C) qPCR assessment of *G6P1* and *G6P2* transcript level in nymphal tick gut after feeding on clean and *B. burgdorferi*-infected mice (ns, *P* > 0.05; *, *P* < 0.05).

**Figure S7.**
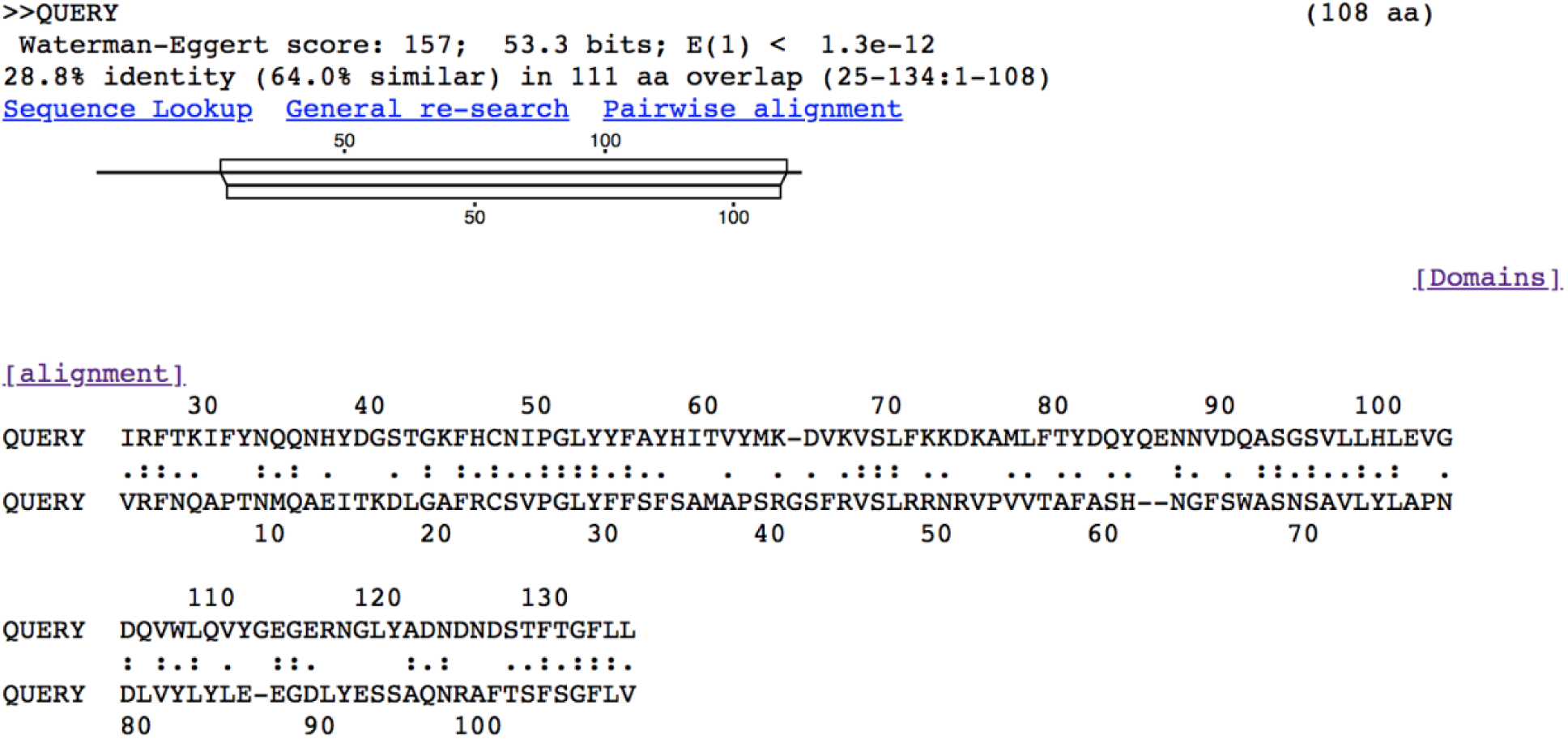
Alignment of human adiponectin C1Q domain and C1QL3 C1Q domain. Tick C1QL3 C1Q domain has 64.0% similarity and 28.8% identity with human adiponectin C1Q domain. The alignment was conducted in LALIGN/PLALIGN (https://fasta.bioch.virginia.edu/fasta_www2/fasta_www.cgi?rm=lalign&pgm=lal).

**Table S1.**
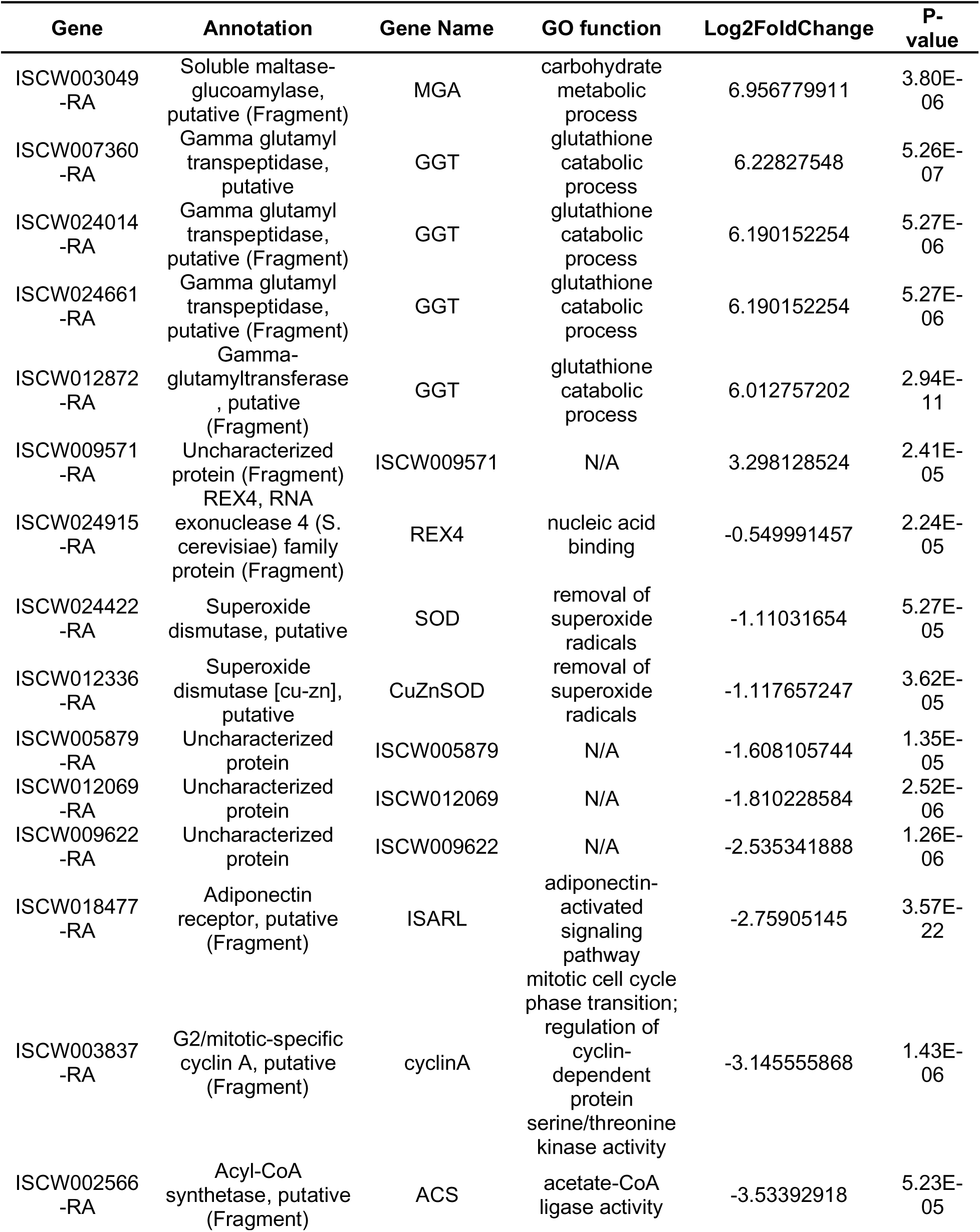
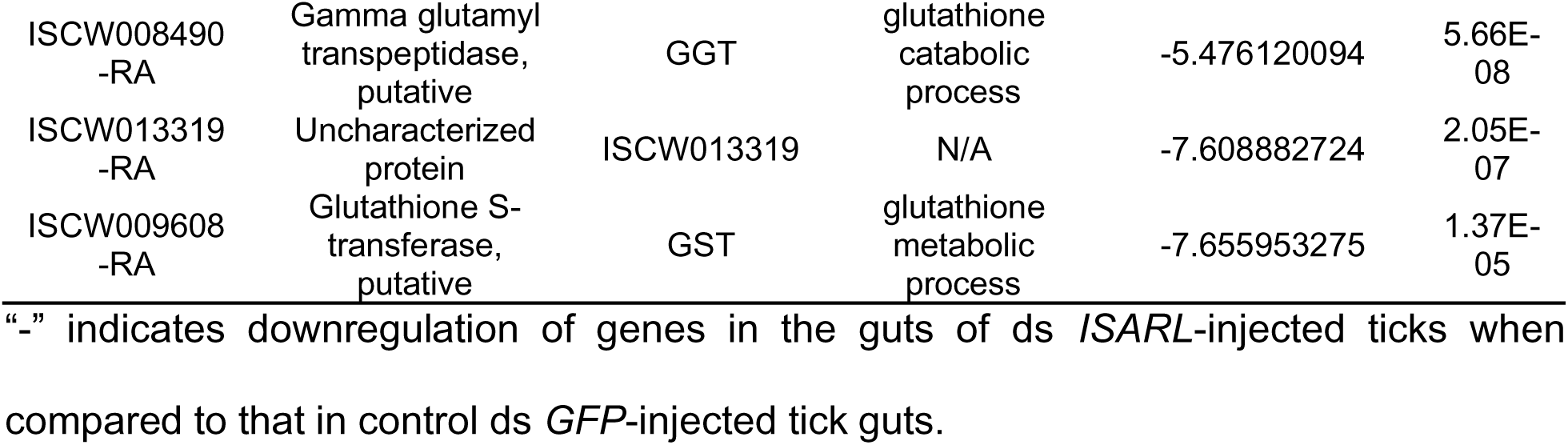
Summary of differently expressed genes of comparison between ds GFP and ds ISARL injection after 96h feeding on clean mice.

**Table S2.** Summary of differently expressed genes of comparison between ds GFP and ds ISARL injection after 96h feeding on B. burgdorferi-infected mice.

| Gene | Annotation | Gene Name | GO function | Log2FoldChange | P-value |
| --- | --- | --- | --- | --- | --- |
| ISCW003218-RA | FADdependent oxidoreductase domain-containing protein 2, putative | FOXRED2 | ubiquitin-dependent ERAD pathway | -2.879056472 | 0.000239 |
| ISCW005667-RA | Secreted protein, putative | Reeler | N/A | -2.678543731 | 0.000119 |
| ISCW003629-RA | Secreted protein, putative | ISCW003629 | N/A | -2.426582135 | 0.0001 |
| ISCW018477-RA | Adiponectin receptor, putative (Fragment) | ISARL | adiponectin-activated signaling pathway | -1.912092255 | 1.72E-11 |
| ISCW011292-RA | Cyclic nucleotidebinding domain-containing protein | CNBD | potassium ion transmembrane transport | -1.620397316 | 0.00021 |
| ISCW003135-RA | Cytochrome p450, putative | CYP | heme binding; oxidoreductase activity | -1.427275425 | 7.35E-05 |
| ISCW001955-RA | AMP dependent CoA ligase, putative | AMP | oxidoreductase activity | -1.427082757 | 0.00027 |
| ISCW002185-RA | Uncharacterized protein | ISCW002185 | N/A | -1.251407193 | 3.40E-05 |
| ISCW024631-RA | 3-hydroxyacyl-CoA dehydrogenase, putative (Secreted salivary gland peptide, putative) | 3HADH | fatty acid metabolic process | -1.089854316 | 2.66E-05 |
| ISCW013566-RA | Alpha-actinin, putative | ACTN | calcium ion binding; protein tyrosine phosphatase activity | -1.051823169 | 4.99E-05 |
| ISCW020505-RA | Uncharacterized protein | ISCW020505 | oxidoreductase activity | -1.038498406 | 4.11E-05 |
| ISCW019656-RA | Secreted salivary gland peptide, putative (Fragment) | ISCW019656 | N/A | -1.000151879 | 5.89E-05 |
| ISCW016391-RA | Cytochrome P450, putative | CYP | heme binding; oxidoreductase activity | -0.983747879 | 0.000348 |
| ISCW006151-RA | Transport protein, putative (Fragment) | ISCW006151 | transmembrane transporter activity | -0.869973215 | 1.75E-05 |
| ISCW021203-RA | Uncharacterized protein | ISCW021203 | N/A | -0.826491344 | 9.19E-05 |
| ISCW024161-RA | Uncharacterized protein | ISCW024161 | N/A | -0.753881171 | 4.99E-05 |
| ISCW018609-RA | Hemomucin, putative (Fragment) | Hmu | biosynthetic process | -0.730567529 | 0.000361 |
| ISCW017282-RA | Oxodicarboxylate carrier protein, putative | ODC | transmembrane transport | -0.701243498 | 4.69E-05 |
| ISCW009471-RA | Phospholipase B-like (Fragment) | PLB | phospholipid catabolic process | -0.694463465 | 8.01E-05 |
| ISCW006150-RA | Sugar transporter, putative | ISCW006150 | carbohydrate transport | -0.666272429 | 6.70E-05 |
| ISCW012299-RA | Adenylosuccinate synthetase (Fragment) | ADSS | de novo' AMP biosynthetic process; IMP metabolic process | -0.649923076 | 0.00038 |
| ISCW022913-RA | cAMP and cAMP-inhibited cGMP 3,5-cyclic phosphodiesterase, putative | PDE | negative regulation of cGMP-mediated signaling | -0.612276289 | 3.11E-06 |
| ISCW000885-RA | Alpha-1,4 glucan phosphorylase | GP | glycogen catabolic process | -0.588614232 | 0.000254 |
| ISCW019117-RA | Pyridoxine kinase, putative (Fragment) | PDXK | pyridoxal 5'-phosphate salvage | -0.535676385 | 0.000478 |
| ISCW018543-RA | GMP synthase, putative (Fragment) | GMPS | glutamine metabolic process; GMP biosynthetic process | -0.53194689 | 0.000465 |
| ISCW004028-RA | Phosphatidylserine synthase I, putative | PTDSS1 | phosphatidylserine biosynthetic process | -0.529598437 | 3.12E-05 |
| ISCW012521-RA | Uncharacterized protein | ISCW012521 | N/A | -0.517853294 | 2.92E-05 |
| ISCW009548-RA | Uncharacterized protein | ISCW009548 | N/A | -0.504914087 | 0.000199 |
| ISCW015732-RA | RNA polymerase II transcription elongation factor, putative | Elongin-C | ubiquitin-dependent protein catabolic process | -0.48942716 | 4.89E-05 |
| ISCW022144-RA | N-CAM Ig domain-containing protein, putative | Ncam | protein serine phosphatase activity; protein threonine phosphatase activity | -0.437192969 | 0.000289 |
| ISCW009549<br>-RA | Selenoprotein P<br>precursor, putative | Selenop | selenium<br>compound<br>metabolic<br>process | -0.436460074 | 0.000423 |
| ISCW013692<br>-RA | V-type proton<br>ATPase subunit G | V-ATPase | proton<br>transmembrane<br>transport | -0.414399839 | 0.000311 |
| ISCW010157<br>-RA | Receptor<br>expression-<br>enhancing protein | REEP | N/A | -0.405156446 | 0.000104 |
| ISCW023312<br>-RA | Sidoreflexin | SFXN | mitochondrial<br>transmembrane<br>transport;<br>serine import<br>into<br>mitochondrion | -0.3999554 | 0.000229 |
| ISCW006433<br>-RA | AP complex subunit<br>sigma (Fragment) | APS | intracellular<br>protein<br>transport;<br>vesicle-<br>mediated<br>transport | -0.395796427 | 0.000508 |

“-” indicates downregulation of genes in the guts of ds *ISARL*-injected ticks when compared to that in control ds *GFP*-injected tick guts.

**Table S3.** Summary of differently expressed genes of comparison between recombinant *GFP* and adiponectin proteins injection after 8h.

| **Gene** | **Annotation** | **Log2FoldChange** | **P-value** |
| --- | --- | --- | --- |
| ISCW004553-RA | Cuticle protein, putative | 18.51302808 | 2.05E-12 |
| ISCW002039-RA | Cuticle protein, putative | 17.49563794 | 2.29E-07 |
| ISCW001782-RA | Uncharacterized protein | 17.27754175 | 7.09E-10 |
| ISCW013798-RA | Cuticle protein, putative | 17.20315541 | 7.96E-05 |
| ISCW015495-RA | Secreted protein, putative (Fragment) | 17.15463748 | 0.00016026 |
| ISCW005191-RA | Secreted glycine rich protein, putative (Fragment) | 17.11093994 | 2.95E-08 |
| ISCW003789-RA | Uncharacterized protein | 16.81877212 | 2.70E-06 |
| ISCW016297-RA | Uncharacterized protein | 16.74612219 | 1.13E-06 |
| ISCW008562-RA | Cuticle protein, putative | 15.3237752 | 1.02E-06 |
| ISCW002925-RA | Uncharacterized protein | 2.857394251 | 1.82E-05 |
| ISCW024478-RA | Secreted cysteine rich protein, putative (Fragment) | 2.851254132 | 1.38E-05 |
| ISCW021558-RA | Secreted salivary gland peptide, putative | 2.689268024 | 3.51E-05 |
| ISCW021555-RA | Uncharacterized protein | 2.656702948 | 4.30E-05 |
| ISCW023547-RA | Secreted salivary gland peptide, putative | 2.589439049 | 0.0001213 |
| ISCW023623-RA | Serpin-4 precursor, putative (Serpin-4, putative) | 2.173997552 | 5.42E-05 |
| ISCW008209-RA | Hebreain, putative | 2.158930697 | 1.20E-06 |
| ISCW024733-RA | Beat protein, putative (Fragment) | 2.105356324 | 1.12E-05 |
| ISCW024387-RA | Serpin-2 precursor, putative (Fragment) | 2.00955178 | 5.56E-05 |
| ISCW002113-RA | Antimicrobial peptide microplusin | 1.940836315 | 0.00014064 |
| ISCW011893-RA | ANK_REP_REGION domain-containing protein | 1.808330643 | 1.28E-08 |
| ISCW015113-RA | Uncharacterized protein | 1.783621042 | 6.95E-05 |
| ISCW012685-RA | Myosin light chain 1, putative | 1.61605594 | 9.51E-05 |
| ISCW005837-RA | Uncharacterized protein | 1.582477028 | 3.43E-05 |
| ISCW024686-RA | Ixoderin, putative (Fragment) | 1.545371762 | 3.67E-06 |
| ISCW014652-RA | Serpin-8 precursor, putative | 1.431124914 | 1.89E-06 |
| ISCW009063-RA | Tropomyosin, putative | 1.358977546 | 0.00014404 |
| ISCW023442-RA | Uncharacterized protein | 1.288696108 | 0.00015793 |
| ISCW023441-RA | Troponin, putative (Fragment) | 1.2703018 | 0.00010777 |
| ISCW016762-RA | LIM domain-containing protein, putative | 1.268776381 | 6.91E-06 |
| ISCW002637-RA | Reductase, putative | 1.195839092 | 5.26E-05 |
| ISCW017459-RA | Glucose-6-phosphatase | -1.182920138 | 0.00012321 |
| ISCW015064-RA | Cytochrome P450, putative | -1.276584666 | 5.75E-06 |
| ISCW015956-RA | Serine/threonine protein kinase, putative | -1.438675023 | 1.25E-05 |
| ISCW016390-RA | Cytochrome P450, putative | -1.643963635 | 3.32E-06 |
| ISCW016391-RA | Cytochrome P450, putative | -1.678990788 | 6.27E-05 |
| ISCW006560-RA | Cytochrome P450, putative | -2.209859052 | 1.42E-06 |
| ISCW024348-RA | Salivary HBP family protein, putative (Fragment) | -3.822772331 | 2.76E-05 |
| ISCW008563-RA | Cuticle protein, putative | -14.56782308 | 1.01E-06 |
| ISCW005940-RA | Elongation of very long chain fatty acids protein | -15.77527062 | 7.81E-08 |
| ISCW012372-RA | Cysteine rich secreted peptide, putative | -17.69968417 | 3.20E-09 |

“-” indicates downregulation of genes in the guts of adiponectin-injected ticks when compared to that in control GFP-injected tick guts.

**Table S4.**
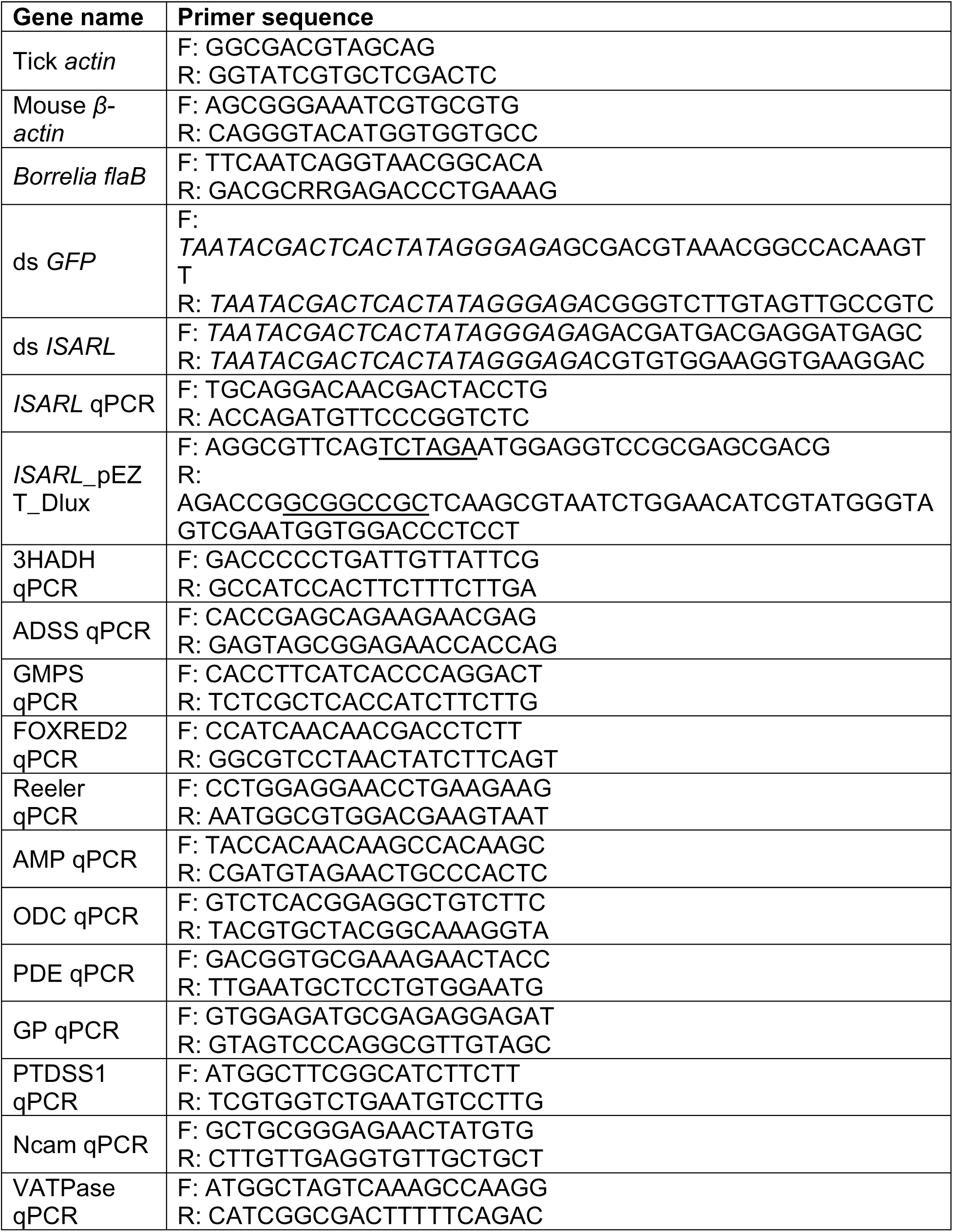
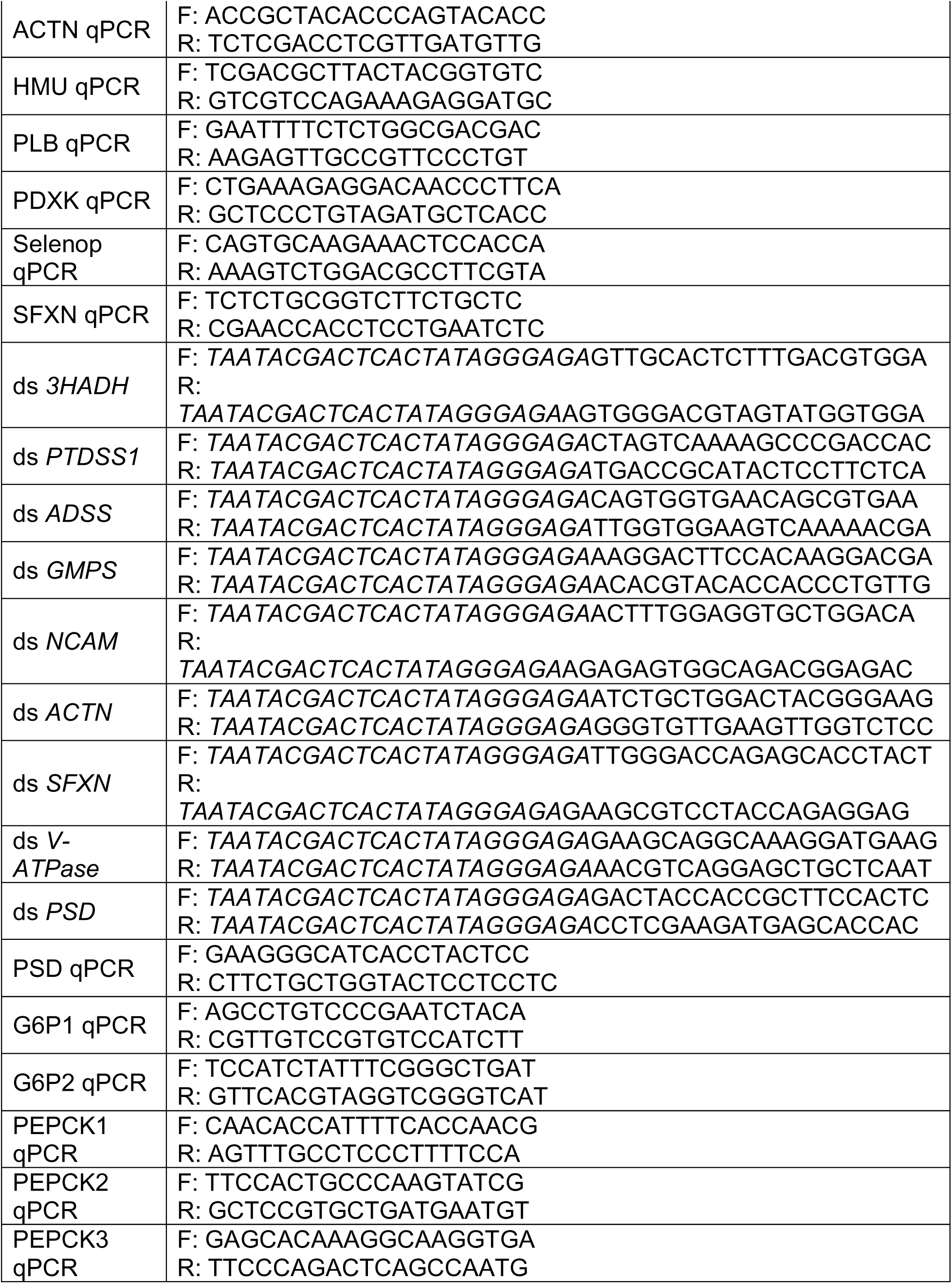
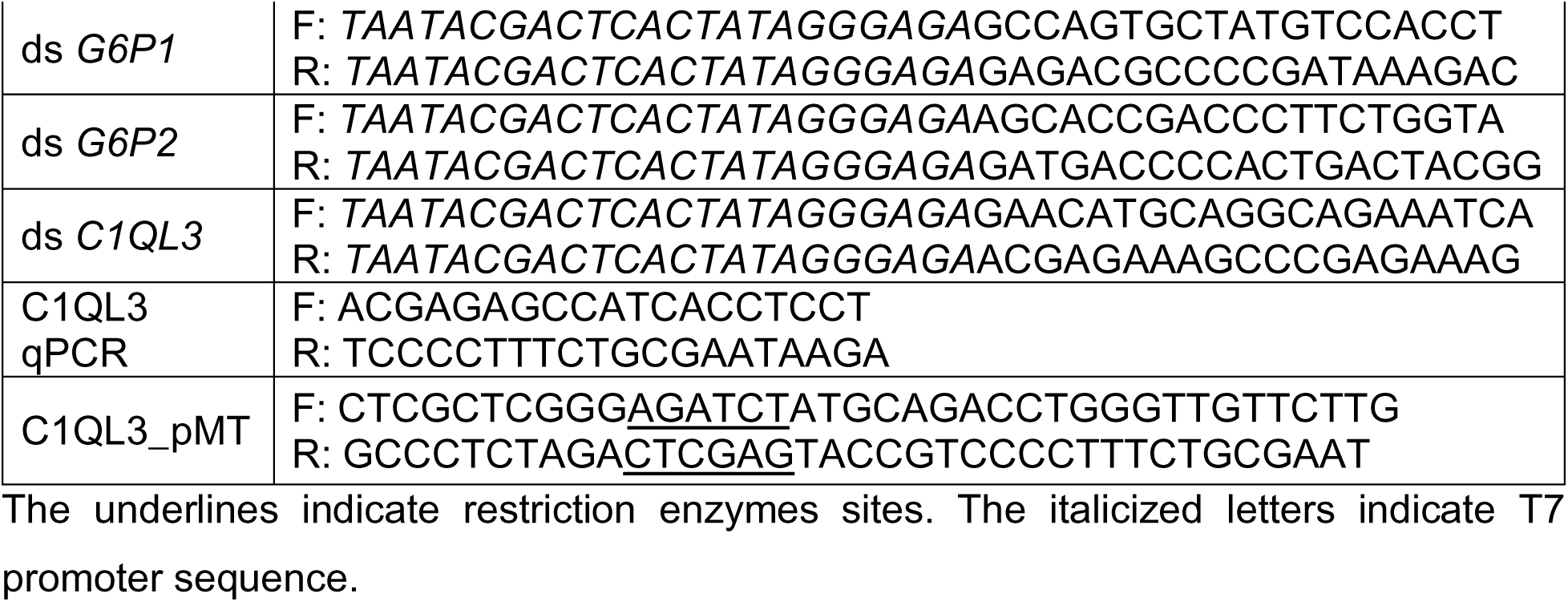
The primers used in this study.

